# Potentially prebiotic synthesis of aminoacyl-RNA via a bridging phosphoramidate-ester intermediate

**DOI:** 10.1101/2022.01.25.477680

**Authors:** Samuel J. Roberts, Ziwei Liu, John D. Sutherland

## Abstract

Translation according to the genetic code is made possible by selectivity both in aminoacylation of tRNA and in anticodon:codon recognition. In extant biology, tRNAs are selectively aminoacylated by enzymes using high-energy intermediates, but how this might have been achieved prior to the advent of protein synthesis has been a largely unanswered question in prebiotic chemistry. We have now elucidated a novel, prebiotically plausible stereoselective aminoacyl-RNA synthesis which starts from RNA-amino acid phosphoramidates and proceeds via phosphoramidate-ester intermediates which subsequently undergo conversion to aminoacyl-esters by mild acid hydrolysis. The chemistry avoids the intermediacy of high-energy mixed carboxy-phosphate anhydrides and is greatly favored under eutectic conditions, which also potentially allow for the requisite pH fluctuation through the variable solubility of CO_2_ in solid/liquid water.

## INTRODUCTION

Ribosomal peptide synthesis relies upon aminoacyl-tRNAs as intermediates in the translation of messenger-RNAs into the corresponding genetically-coded protein products. Highly specific enzymatic aminoacylation of the 2’,3’-diol of tRNAs using aminoacyl-adenylates is crucial to this process. In the absence of enzymes, which bind them tightly and protectively, carbon dioxide promotes the reversible conversion of aminoacyl-adenylates to amino acid *N*-carboxyanhydrides (NCAs) and the latter are subsequently polymerized to random peptides in aqueous solution^1,2^ Other aminoacyl-RNA mixed anhydrides **1** have been used as substrates for non-enzymatic aminoacyl-RNA synthesis in prebiotic model studies^3,4^, but they are similarly susceptible to carbon dioxide promoted destruction. Conceptually, there are two fundamentally different chemical pathways for the formation of mixed anhydrides – via either carboxylate activation or phosphate activation (the latter of which is used by modern day biology). In the former, the phosphate reacts (reversibly) with an activated carboxylate (e.g. NCA, 5(4*H*)-oxazolone or other species) giving a mixed anhydride **1**.^5–7^ Conversely, in the phosphate activation pathway a native amino acid reacts with an activated phosphate (e.g. imidoyl phosphate) giving a mixed anhydride **1**.^7^ Carboxylate activation processes^5,6^ are inherently problematic under a carbon dioxide containing atmosphere in that the formation of NCAs and hence random non-coded peptides cannot be avoided.

Prebiotic activation chemistry is also needed to enable the formation of phosphodiesters from alcohols and the corresponding phosphomonoesters. Thus, for example, 5’-phosphorimidazolides are used for prebiotic RNA ligation or monomer extension chemistry^8,9^. Seeking to unify the various building block activation chemistries, we were intrigued by the work of Orgel *et al*. showing that amino acids can react with 5’-phosphorimidazolides through their amino groups generating amino acid phosphoramidates **2** (Scheme 1)^10^. By using adenine nucleotide 2’/3’-aminoacyl-esters and 5’-phosphorimidazolides, bridged phosphoramidate-esters could be formed by templating with poly(U)^11^. The Richert group has built upon these earlier results and shown that RNA-peptide phosphoramidates can couple to 3’-amino-2’,3’-dideoxy-terminated oligonucleotides to give bridging peptide phosphoramidate-amides on an RNA template^12^. However, the reaction of RNA-amino acid phosphoramidates **2** (Scheme 1) with the 2’,3’-diol of a canonical oligonucleotide **3** to generate phosphoramidate-esters **4**, is inherently more interesting as the products are potentially hydrolyzable to aminoacyl-esters **5** and 5’-phosphoryl-RNAs **6** under mildly acidic conditions^13^. Synthesis of aminoacyl-esters in this way would bypass the unstable mixed anhydrides **1** employed in other strategies^14^ and might have been more easily achievable before the advent of enzymes.

We have previously postulated that loops could be provide chemoselectivity in aminoacylation and avoid the necessity of extra strands of RNA^14^. tRNAs have 5’-phosphate groups and a short 3’-overhang at the end of the acceptor stem and accordingly we wondered if phosphoramidate-ester **4** formation would be possible in such a stem-overhang configuration. We chose UUCCA as an overhang sequence as it shows strong sequence similarity to the CCA overhang of modern tRNAs and was a successful sequence identified in our previous work^14^.

## RESULTS

A chemically synthesized RNA-amino acid phosphoramidate (**2-Gly**, 3’-AGCGAp-Gly-OH, 100 μM) was mixed with ester acceptor RNA strand **3** (3’-ACCUUUCGCU, 100 μM), and activation buffer (made up of the water-soluble carbodiimide EDC and imidazole in buffered aqueous solution). Whilst EDC is prebiotically implausible we use it here as others have done previously as a model activating agent allowing formation of an acyl imidazolide^12^. The reaction was incubated at room temperature overnight and afterwards analyzed by HPLC (Figure SI-1). In comparison to controls containing 5’-phosphoryl-RNA **6** in place of **2-Gly**, (Figure SI-17), an extra peak could be observed in the HPLC trace of the reaction products consistent with the formation of phosphoramidate-ester **4-Gly**. After purification by HPLC, this product was shown by MALDI-TOF mass spectroscopy to indeed have the mass of **4-Gly** (3’AGCGAp-Gly-ACCUUUCGCU, calculated M.W. 4767.7, found: 4768.0 [M+H]). This peak also disappeared on treatment with alkali, consistent with the presence of an ester linkage in the product^15^.

**Scheme 1.**
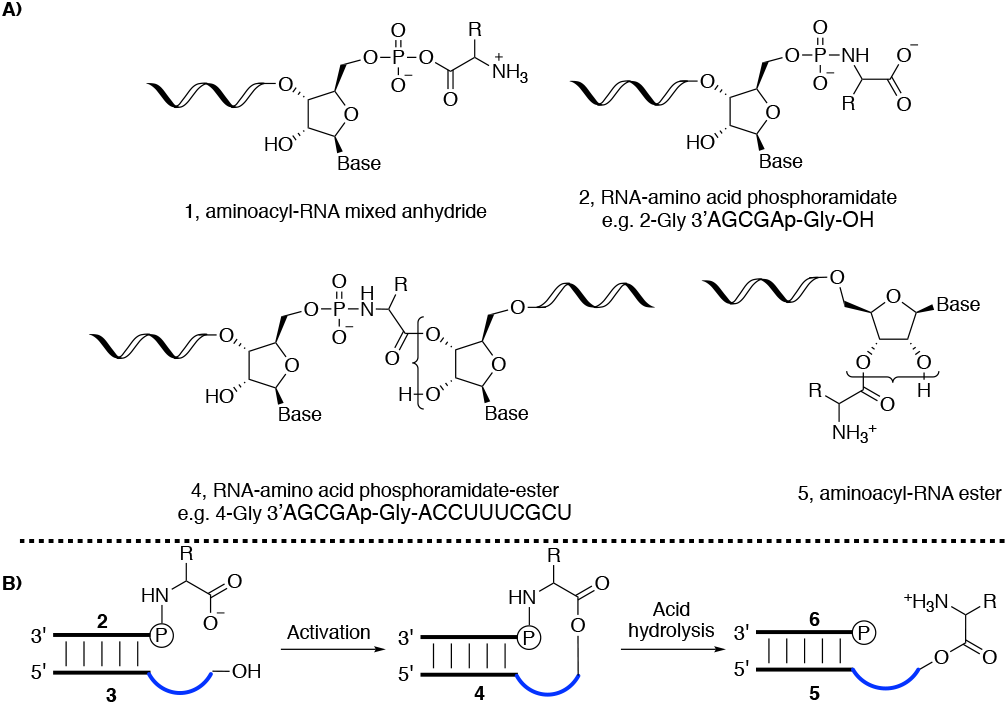
**(a) Chemical structures of RNA-amino acid hybrid compounds. (b) Reaction scheme for the formation of aminoacyl-RNA esters. RNA-amino acid phosphoramidate-ester (middle structure, 4) is formed from RNA phosphoramidate (2, left) and ester acceptor RNA (3) under activation conditions. The phosphoramidate bond can then be cleaved under mild acid conditions generating aminoacyl-RNA (5, right)**.

Encouraged by these findings, we further synthesized two different RNA-amino acid phosphoramidates (**2-L-Ala**, 3’AGCGAp-L-Ala-OH and **2-D-Ala**, 3’AGCGAp-D-Ala-OH) to explore their comparable reactivity and the stereoselectivity of ester formation. Based on ^31^P-NMR spectra (Figure SI-19), the yields of formation of **2-L-Ala** and **2-D-Ala** from their parent 5’-phosphoryl-RNAs were similar, suggesting that there is no stereoselectivity in the conventional synthesis of phosphoramidates of single stranded RNA. After HPLC purification, each of the RNA-amino acid phosphoramidates was combined with 10-mer RNA ester acceptor **3** (3’-ACCUUUCGCU, 100 μM) in activation buffer and again new HPLC peaks were evident after reaction. As shown in Figures SI-2 and SI-3, the L- and D-alanine phosphoramidate-esters **4-L-Ala** and **4-D-Ala** were formed after 18 hours, their identity confirmed by alkaline hydrolysis and the former additionally by MALDI-TOF mass spectra (**4-L-Ala**: 3’AGCGAp-L-Ala-ACCUUUCGCU, calculated M.W. 4781.7, found: 4782.2 [M+H]). Interestingly, the yield of **4-D-Ala** was significantly lower than that of **4-L-Ala** (9.5:1, quantified as an average of the ratios across three replicates using an internal standard). It is known that aminoacyl-imidazolides are key to the aminoacylation of one of the hydroxyl groups of a diol in water^16^, presumed due to the protonation of the imidazole leaving group mediated through the second hydroxyl group^2^, and we suspect that acyl-imidazolides are similarly crucial to the chemistry we have discovered as we observed no phosphoramidate ester **4-L-Ala** product when using activation buffer in the absence of imidazole (Figure SI-14). Other non-base sensitive peaks also appeared independent of the presence of imidazole. It has been reported that EDC (albeit at much higher concentrations) reacts with RNA to form cyclic structures or modify nucleobases, and this could account for some of the other base insensitive peaks we observe (Figure SI-1 to SI-18)^17, 18^.

With these results in hand, we next wanted to scope out the chemoselectivity and stereoselectivity for different amino acids. Using the same oligonucleotide sequence as before, we isolated phosphoramidates **2** of leucine, valine, arginine, proline and serine covering a range of prebiotically plausible early amino acids^19^. On individually submitting all 5 RNA-L-amino acid phosphoramidates to ester forming conditions in activation buffer with 10-mer RNA ester acceptor **3** (3’-ACCUUUCGCU, 100 μM), peaks for all phosphoramidate-ester products could be identified (correct MALDI-TOF masses and susceptibility to alkaline hydrolysis) (Figures SI-4 to SI-11, Table SI-1). However, the yields varied according to the amino acid side chain (Figure 1). Comparable reactions using D-amino acid phosphoramidates all afforded significantly lower yields of phosphoramidate-esters compared to the corresponding L-amino acid phosphoramidates (Table SI-3). Thus it appears that with the esterification chemistry described herein there is a general relative stereochemical correlation between L-amino acids and RNA based on D-ribose.

**Figure 1.**
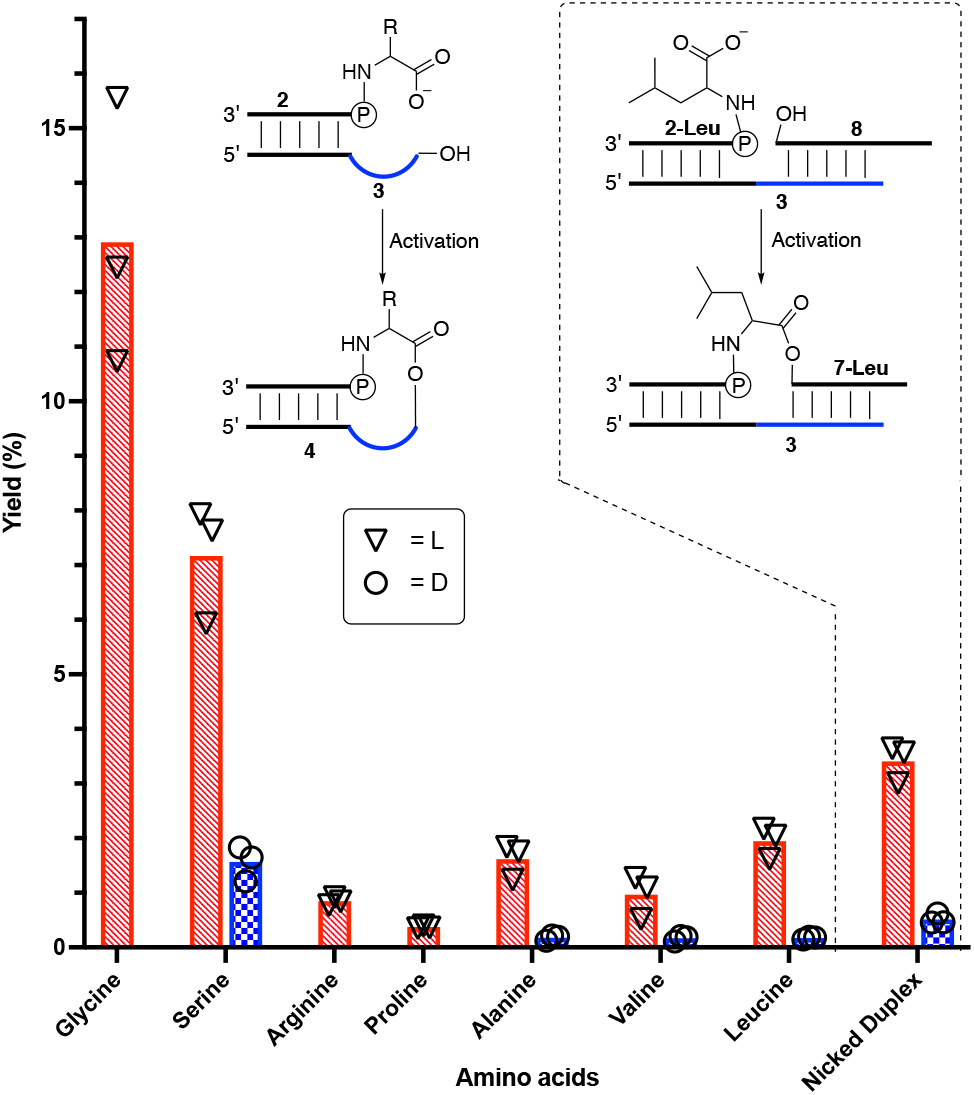
Room temperature yields for formation of phosphoramidate-esters **4** via a nicked loop and **7-Leu** via a nicked duplex. Bars are mean values based on three replicates. Each replicate is represented by a circular/triangular data point. Red hashed bars represent L-amino acid stereochemistry and blue checkered bars D- (where appropriate).

In some cases, HPLC peaks corresponding to the phosphoramidate-ester products **4** were observed with ‘shoulders’ or as double peaks (e.g. Figure SI-4). In all cases we found that both peaks decreased in intensity upon alkaline hydrolysis although not always at the same rate. These observations suggest the formation of mixtures of 2’- and 3’-linked phosphoramidate-esters **4**. Similar aminoacyl-migration is known with aminoacyl-tRNAs^20^ and so we have not concerned ourselves further with this aspect of regioselectivity.

High stereoselectivity and at least moderate chemoselectivity for specific amino acids would be features essential for the development of genetic encoding of biologically useful peptides before the emergence of enzymatic aminoacylation. We have previously established that variations in RNA sequence are associated with different yields in another aminoacyl-transfer system.^14^

It is possible that in sequences that contain a palindromic (or partially palindromic) sequence the esterification could occur through a nicked-duplex type configuration rather than by way of a stem with a folded back overhang. Alternatively, complimentary sequences of RNA may bind to the overhang sequence, producing a nicked-duplex. To study this we decided to impose a nicked-duplex configuration in our system by including an additional oligonucleotide with a sequence complementary to the loop region (Figure 1). The reaction of amidate **2-L-Leu**, template **3** and complement 3’-AAGGUAAU **8** in activation buffer produced a new alkali sensitive peak with a long HPLC retention time, whilst no trace of the previously observed phosphoramidate-ester product **4-L-Leu** could be detected. This new peak was isolated and shown to have a mass consistent with the 13mer phosphoramidate-ester **7-L-Leu** (3’AGCGAp-Leu-AAGGUAAU, calculated M.W. 4323.7, found: 4324.2 [M+H]) and to be alkali sensitive. On repeating the experiment using the D-leucine phosphoramidate **2-D-Leu**, we saw a greatly reduced intensity peak posited, on the basis of alkaline sensitivity, to be due to the D-Leu-13mer **7-D-Leu** (L:D 7:1 compared to L:D 12:1 for leucine nicked-loop esterification, Figure SI-12 to SI-13). This result suggests that any sequences which allow esterification via a nicked-duplex configuration will have reduced L:D selectivity.

Serendipitously, on resubmitting a sample to HPLC analysis that had been stored at -16 °C for a week after the initial reaction at room temperature, we observed greatly increased intensity of the phosphoramidate-ester product peak. However, samples kept at -70 °C showed no evidence of similarly enhanced reaction. This suggested that the esterification reaction could take place under eutectic conditions with increased yields. After submitting fresh samples to -16 °C in activation buffer for 7 days, the L-amino acid phosphoramidate-ester products formed in much higher yields (Figure 2), whilst formation of D-amino acid phosphoramidate-esters did not improve significantly over the room temperature yields, resulting in improved L/D selectivity over the room temperature conditions (Table SI-3). Yields increased further by 14 days, but the L:D selectivity then decreased, likely due to lower levels of starting materials available in the ‘L-reactions’ over the second week. As the individual data points of Figure 2 show, there is a large variation between replicates of the eutectic samples despite the high reproducibility of results from the room temperature reaction (Figure 1) from the same starting mixture. We suggest that, amongst other things, this could be due to relatively poor temperature control in a large -16 °C freezer.

**Figure 2.**
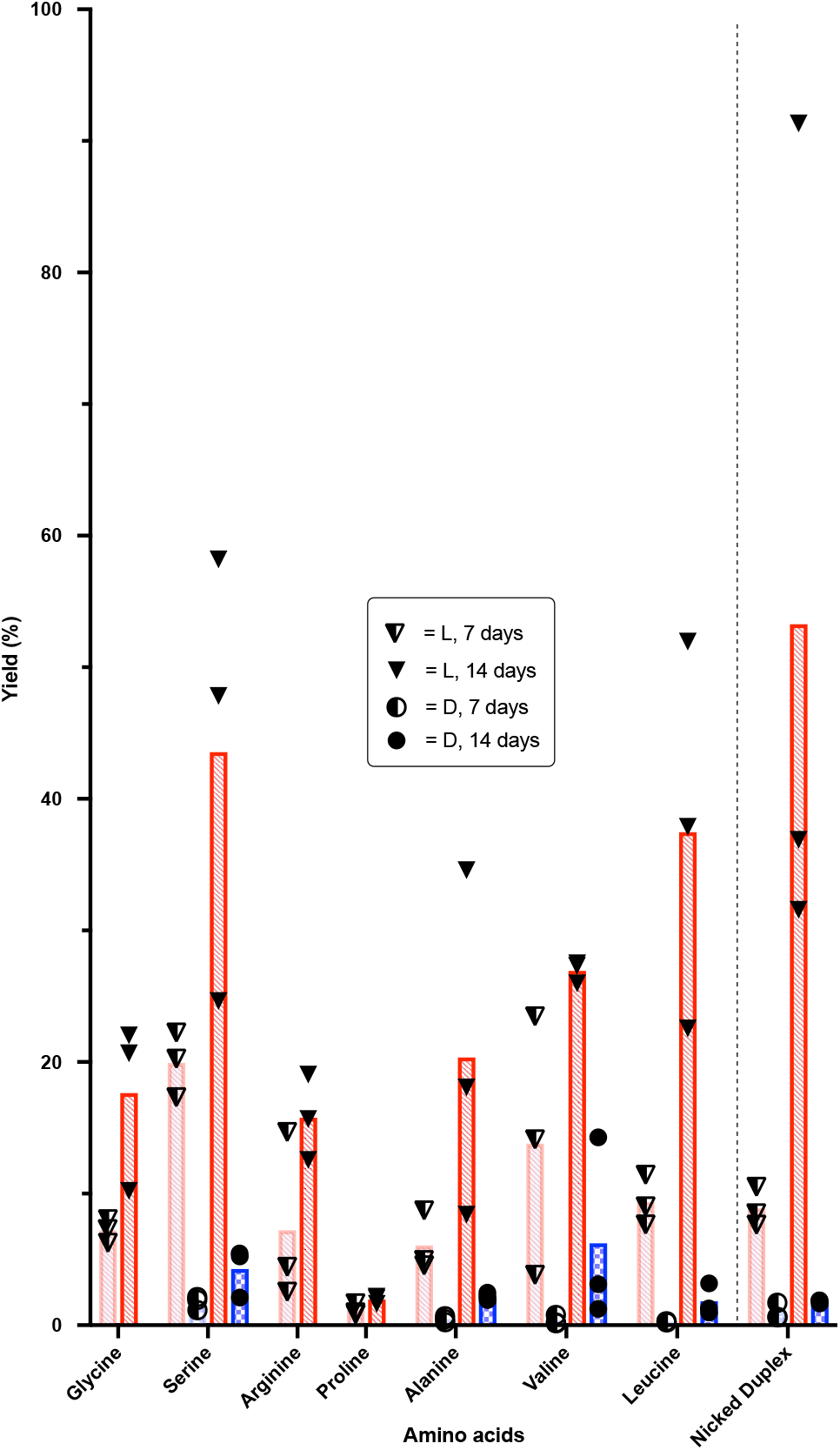
Eutectic yields for formation of phosphoramidate-esters **4** via a nicked loop and **7-Leu** via a nicked duplex. Each replicate is represented by a circular/triangular data point. Red hashed bars represent L-amino acid stereochemistry and blue checkered bars D- (where appropriate). Darker colors indicate longer reaction times.

We next studied the formation of aminoacyl-RNA esters **5** by mild acid hydrolysis of phosphoramidate-esters **4** (Figure 3). The HPLC purified phosphoramidate-esters **4** were dissolved in pH = 3 formate buffer (final concentration 83 mM formate) with an internal HPLC standard and kept at 25 °C overnight. HPLC analysis revealed that all the phosphoramidate-ester **4** was hydrolyzed, giving 5’-phosphoryl-5-mer **6** RNA (3’-AGCGAp), aminoacyl-10mer **5** (H-amino acid-3’-ACCUUUCGCU) and 10-mer RNA **3** (3’-ACCUUUCGCU), which resulted from subsequent slow ester hydrolysis. The free aminoacyl-ester **5** was produced quantitatively or near quantitatively for glycine, alanine, valine and leucine (the latter either from the product of the nicked-loop or nicked-duplex chemistry), with little difference observed between L- and D-valine phosphoramidate esters **4-L-Val** and **4-D-Val** (the only amino acid for which we could isolate sufficient D-phosphoramidate ester to study). The identity of 10-mer RNA **3** was confirmed by spiking with an authentic sample and the identity of aminoacyl-RNA **5** was confirmed by its susceptibility to alkaline hydrolysis. After such hydrolysis, the aminoacyl-RNA **5** peak disappeared, and the peak corresponding to the 10-mer RNA **3** increased adding further weight to the characterization (Figure SI-21 to SI-34). To varying degrees with different amino acids we also identified 5’-hydroxyl-5mer **9** (e.g. Figure SI-22). We hypothesize that this likely resulted from hydrolysis of the ester bond of the phosphoramidate ester **4** (to produce 10-mer RNA **3** and phosphoramidate-5mer **2**), followed by internal hydrolysis either in a similar manner to that previously hypothesized by the McGuigan group in relation to ‘Protide’ activation^21^, or by in-line hydrolysis to afford **9**. On submitting all amino acid-phosphoramidate starting materials **2** individually to pH = 3 formate buffer (60 mM final concentration formate), we identified only 2 products – 5’-phosphoryl-5-mer **6** and 5’-hydroxyl-5-mer **9** – in varying yields (Table SI-2), with a trend for increased formation of **9** following the Thorpe-Ingold effect for alanine, valine and leucine. What’s more, we also saw a consistent increase in yield of **6** with D-relative to L-amino acids. In all cases, acidification of phosphoramidate-esters **4** predominantly resulted in hydrolysis of the phosphoramidate over the ester. With the exception of serine (Figure SI-30) no unassignable peaks were observed for any of acid hydrolyses, implying the conditions maintain the integrity of the RNA.

**Figure 3.**
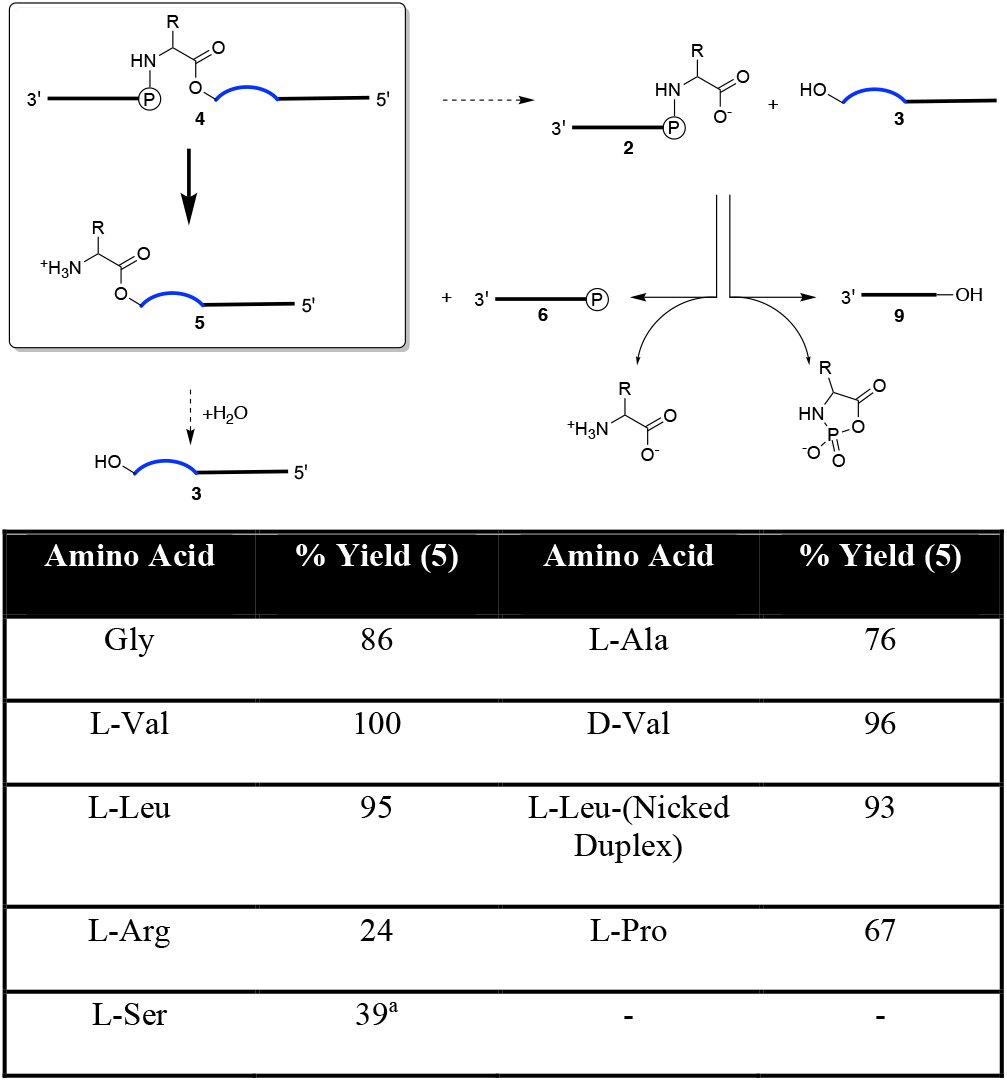
Top - Mechanism of hydrolysis of the phosphoramidate ester **4** at pH = 3. The majority of **4** undergoes phosphoramidate bond hydrolysis giving 5’-phosphoryl-5-mer **6** and the desired aminoacyl-10mer **5** product which can then degrade slowly back to template **3**. Minority hydrolyses at the ester bond of **4** gives **3** and 5’ phosphoramidate-5-mer **2**. Phosphoramidate **2** degrades either by direct hydrolysis to 5’-phosphoryl-5-mer **6**, or by intramolecular oligonucleotide dephosphorylation to the 5’-hydroxyl-5-mer **9** in an amino acid specific fashion. Bottom – Yields of formation of aminoacyl-10mer **5** from phosphoramidate-ester **4** with different amino acids. ^a^. Estimate due to unknown products of hydrolysis. A discussion of the differing yields of Arg, Pro and Ser is presented in the ESI.

## DISCUSSION

This work has demonstrated that efficient production of aminoacyl-RNAs **5** can be achieved via stepwise esterification of RNA-amino acid phosphoramidates **2** to proximal oligonucleotide-2’,3’-diols **3** followed by mild acid hydrolysis of the phosphoramidate linkage. The lack of acidic degradation could be attributed to the short length of treatment (17 hours) a factor which would have to be incorporated into the prebiotic scenario. The proximity required to enable esterification can be realized in nicked-loop or nicked-duplex configurations. The esterification reaction is stereoselective – being preferred for L-over D-amino acid residues with RNA based on D-ribose – and the stereoselectivity is highest in the nicked-loop configuration. The reaction with one set of oligonucleotide sequences exhibits some chemoselectivity for different amino acid side chains auguring well for higher chemoselectivity when sequence variability is combinatorially explored in future studies. If this chemistry were realised with glutamic or aspartic acids it is likely that the prebiotic activating agent would also activate their side chains to nucleophilic attack. Whether these amino acids are relevant at the origins of translation would have to take this into account.

Yields of the intermediate phosphoramidate ester **4** were greatly improved when the reaction was performed under eutectic conditions as was stereoselectivity, but we do not know why. The rates of ester formation and hydrolysis and competing direct EDC hydration are all likely to be slowed at -16 °C, but probably differentially. The lower temperatures are also likely to promote more rigid RNA secondary structure and this could enhance stereoselectivity. The effects of salt concentration and pH are well documented in the eutectic phase^22,23^, but there are also likely to be other effects at play that we have not considered and whilst eutectic effects on RNA chemistry have been studied at length^24,25^, understanding the effect on these conditions in more detail is clearly required.

The second reaction in the sequence occurs under different conditions to the first and it is tempting to speculate about environmental effects that could cause this change. Carbon dioxide is substantially less soluble in ice than it is in water and this, along with the previously documented partitioning of HCl from brine into ice^26^, could contribute to the pH under eutectic conditions being elevated relative to the fully thawed state. The dilution and acidification accompanying melting of icy brines under a carbon dioxide rich atmosphere could favor both the phosphoramidate bond hydrolysis required to generate aminoacyl-esters and enable RNA duplex dissociation^27^. Upon refreezing, the pH of the sample would increase as would the concentration of solutes and RNA leading to the re-adoption of duplex and other structures. If the aminoacyl-esters of two different stem-overhangs were thereby brought into proximity, transpeptidation might ensue. Studies to investigate this possibility with a view to understanding the origin of (coded) peptide synthesis are now underway.

## ASSOCIATED CONTENT

### Supporting Information

Experimental methods including RNA synthesis methods, and HPLC chromatograms and additional data are include in the supporting information. This material is available free of charge via the Internet at http://pubs.acs.org.”

## ACKNOWLEDGMENT

The authors thank Aleksandar Radakovic, Long-Fei Wu and Jack W. Szostak and other J. D. S. group members for fruitful discussions.

---

SYNOPSIS TOC (Word Style “SN_Synopsis_TOC”)

If you are submitting your paper to a journal that requires a synopsis graphic and/or synopsis paragraph, see the Instructions for Authors on the journal’s homepage for a description of what needs to be provided and for the size requirements of the artwork.

To format double-column figures, schemes, charts, and tables, use the following instructions:

> Place the insertion point where you want to change the number of columns
>
> From the **Insert** menu, choose **Break**
>
> Under **Sections**, choose **Continuous**
>
> Make sure the insertion point is in the new section. From the **Format** menu, choose **Columns**
>
> In the **Number of Columns** box, type **1**
>
> Choose the **OK** button

Now your page is set up so that figures, schemes, charts, and tables can span two columns. These must appear at the top of the page. Be sure to add another section break after the table and change it back to two columns with a spacing of 0.33 in.

**Table 1.**
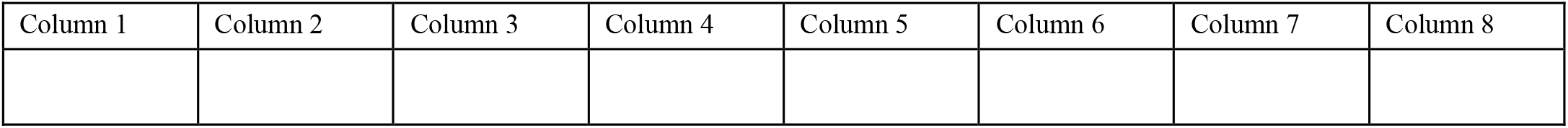
Example of a Double-Column Table.

Authors are required to submit a graphic entry for the Table of Contents (TOC) that, in conjunction with the manuscript title, should give the reader a representative idea of one of the following: A key structure, reaction, equation, concept, or theorem, etc., that is discussed in the manuscript. Consult the journal’s Instructions for Authors for TOC graphic specifications.

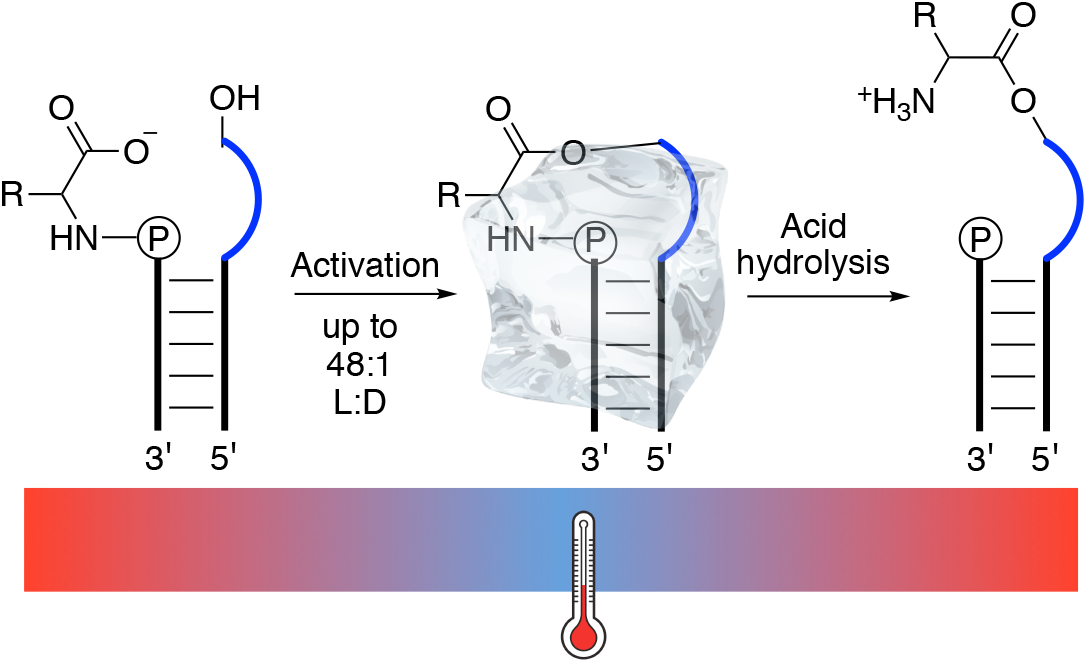

## REFERENCES

(1) Liu, Z.; Beaufils, D.; Rossi, J.C.; Pascal, R., Evolutionary importance of the intramolecular pathways of hydrolysis of phosphate ester mixed anhydrides with amino acids and peptides. Scientific reports, 2014, 4(1), 1–7.

(2) Liu, Z.; Hanson, C.; Ajram, G.; Boiteau, L.; Rossi, J.C.; Danger, G.; Pascal, R., 5 (4H)-Oxazolones as effective aminoacylation reagents for the 3′-terminus of RNA. Synlett, 2017, 28(01), 73–77

(3) Tamura, K.; Schimmel, P. Chiral-selective aminoacylation of an RNA minihelix. Science 2004, 305, 1253.

(4) Tamura, K.; Schimmel, P. Chiral-selective aminoacylation of an RNA minihelix: Mechanistic features and chiral suppression. Proc. Natl. Acad. Sci. U. S. A. 2006, 103, 13750−13752.

(5) Leman, L.; Orgel, L.; Ghadiri, M.R. 2004. Carbonyl sulfide-mediated prebiotic formation of peptides. Science, 2004, 306(5694), 283–286.

(6) Leman, L. J.; Orgel, L. E.; Ghadiri, M. R. Amino acid dependent formation of phosphate anhydrides in water mediated by carbonyl sulfide. J. Am. Chem. Soc. 2006, 128, 20−21.

(7) Liu, Z.; Wu, L.-F.; Xu, J.; Bonfio, C.; Russell, D. A.; Sutherland, J. D. Harnessing chemical energy for the activation and joining of prebiotic building blocks. Nat. Chem. 2020, 12, 1023−1028.

(8) Weimann, B.J.; Lohrmann, R.; Orgel, L.E.; Schneider-Bernloehr, H.; Sulston, J.E. Template-directed synthesis with adenosine-5′-phosphorimidazolide. Science 1968, 161, 387;

(9) Walton, T.; Szostak, J.W., A highly reactive imidazolium-bridged dinucleotide intermediate in nonenzymatic RNA primer extension. J. Am. Chem. Soc. 2016, 138, 11996–12002

(10) Weber, A.L.; Orgel, L.E., Amino acid activation with adenosine 5′-phosphorimidazolide. J. Mol. Evol. 1978, 11, 9–16

(11) Shim, J.L.; Lohrmann, R.; Orgel, L.E. Poly (U)-directed transamidation between adenosine 5’-phosphorimidazolide and 5’-phosphoadenosine 2’(3’)-glycine ester. J. Am. Chem. Soc., 1974, 96, 5283–5284.

(12) Jash, B.; Richert, C. Templates direct the sequence-specific anchoring of the C-terminus of peptido RNAs. Chem. Sci., 2020, 11, 3487–3494.

(13) Liu, Z., Ajram, G., Rossi, J.C. and Pascal, R., The chemical likelihood of ribonucleotide-α-amino acid copolymers as players for early stages of evolution. J. Mol. Evol., 2019, 87, 83–92.

(14) Wu, L.F., Su, M., Liu, Z., Bjork, S.J. and Sutherland, J.D., Interstrand Aminoacyl Transfer in a tRNA Acceptor Stem-Overhang Mimic. J. Am. Chem. Soc., 2021, 143, 11836–11842

(15) Radakovic, A., Wright, T.H., Lelyveld, V.S. and Szostak, J.W., A Potential Role for Aminoacylation in Primordial RNA Copying Chemistry. Biochemistry, 2021, 60, 477–488.

(16) Profy, A.T.; Usher, D.A., Stereoselective aminoacylation of a dinucleoside monophosphate by the imidazolides of dl-alanine and N-(tert-butoxycarbonyl)-dl-alanine. Journal of molecular evolution, 1984 20(2), 147–156.

(17) Obianyor, C.; Newnam, G.; Clifton, B. E.; Grover, M. A.; Hud, N. V. Towards Efficient Nonenzymatic DNA Ligation: Comparing Key Parameters for Maximizing Ligation Rates and Yields with Carbodiimide Activation. ChemBioChem, 2020 21(23), 3359–3370.

(18) Edeleva, E.; Salditt, A.; Stamp, J.; Schwintek, P.; Boekhoven, J.; Braun, D. Continuous nonenzymatic cross-replication of DNA strands with in situ activated DNA oligonucleotides. Chem. Sci., 2019 10(22), 5807–5814.

(19) Patel, B. H.; Percivalle, C.; Ritson, D. J.; Duffy, C. D.; Sutherland, J. D. Common origins of RNA, protein and lipid precursors in a cyanosulfidic protometabolism. Nat. Chem. 2015, 7, 301−307.

(20) Griffin, B. E.; Jarman, M.; Reese, C. B.; Sulston, J. E.; Trentham, D. R. Some observations relating to acyl mobility in aminoacyl soluble ribonucleic acids. Biochemistry, 1966 5(11), 3638–3649.

(21) Mehellou, Y.; Balzarini, J.; McGuigan, C., Aryloxy phosphoramidate triesters: a technology for delivering monophosphorylated nucleosides and sugars into cells. ChemMedChem 2009, 4, 1779 – 1791

(22) Vajda, T., Cryo-bioorganic chemistry: molecular interactions at low temperature. Cell. Mol. Life Sci. 1999, 56, 398–414.

(23) Williams-Smith, D.L.; Bray, R.C.; Barber, M.J.; Tsopanakis, A.D.; Vincent, S.P., Changes in apparent pH on freezing aqueous buffer solutions and their relevance to biochemical electron-paramagnetic-resonance spectroscopy. Biochem. J. 1977, 167, 593–600

(24) Attwater, J., Wochner, A., Pinheiro, V.B., Coulson, A. and Holliger, P., Ice as a protocellular medium for RNA replication. Nat. Commun. 2010, 1, 76;

(25) Mutschler, H. and Holliger, P., Non-canonical 3′-5′ extension of RNA with prebiotically plausible ribonucleoside 2′, 3′-cyclic phosphates. J. Am. Chem. Soc. 2014, 136, 5193−5196

(26) Takenaka, N.; Tanaka, M.; Okitsu, K.; Bandow, H. Rise in the pH of an unfrozen solution in ice due to the presence of NaCl and promotion of decomposition of gallic acids owing to a change in the pH. J. Phys. Chem. A 2006, 110, 10628– 10632

(27) Mariani, A., Bonfio, C., Johnson, C.M. and Sutherland, J.D., pH-Driven RNA strand separation under prebiotically plausible conditions. Biochemistry, 2018, 57, 6382–6386

