## Supplementary Information for "Potentially prebiotic synthesis of aminoacyl-RNA via a bridging phosphoramidate-ester intermediate"

Electronic Supplementary Information

Additional comments	2

Comment on yields of Arginine, Serine and Proline phosphoramidate-esters (**4**)	2

Methods	2

General Methods	2

Chemical synthesis of RNA oligomers	3

Chemical synthesis of RNA phosphoramidate (**2**): Method 1	3

Chemical synthesis of RNA phosphoramidate (**2**): Method 2	4

The formation of RNA phosphoramidate-esters (**4**) or (**7**)	4

Phosphoramidate-ester (**4**) purification	5

Acid hydrolysis of phosphoramidate-ester (**4**)	5

Base hydrolysis of phosphoramidate-ester (**4**)	5

Base hydrolysis of acid treated phosphoramidate-ester (**4**)	6

Experimental Data	7

Phosphoramidate-ester (**4**) formation	7

Phosphoramidate (**2**) formation	28

Hydrolyses	30

Additional Data	44

Control for Reaction of Free Amino Acid Under Phosphoramidate Ester Forming Conditions…………………………………………………………………………………………………………………….. 44

Phosphoramidate (**2**) and phosphoramidate-ester (**4**) Mass Spec data	45

Yield of 5'P-5mer (**6**) produced from hydrolysis of RNA phosphoramidate (**2**)	47

HPLC calibration curves	48

Stereoselectivity of phosphoramidate-ester (**4**) formation	51

### Additional comments

##### Comment on yields of Arginine, Serine and Proline phosphoramidate-esters (4)

Based on these results we can make some inferences about why some of the yields of ester **5** from phosphoramidate-ester **4** vary. Arginine amidate **2-L-Arg** nearly entirely converts (94%) to 5'-hydroxyl-5mer **9** at pH 3, however when **4-L-Arg** is submitted to pH 3 we only saw a 35% yield of **9**. Consequently, as we only see 24% yield of ester **5-L-Arg**, the remaining ~39% of material must be 10mer **3** formed from ester hydrolysis of **5**. In contrast, proline amidate **2-L-Pro** only produces 13% **9** (likely due to the steric constraints of its ring) in acid. Similarly the phosphoramidate-ester **4-L-Pro** is also low yielding in **9** (~7%). This suggests that **5-L-Pro** is largely stable to acid as the majority of the production of **3** can be accounted for by ester hydrolysis of **4-L-Pro** and that the lower yield of **5-L-Pro** is entirely due to ester hydrolysis of the starting material **4-L-Pro**. Finally, production of **5-L-Ser** is complicated as other base sensitive by-products are also observed to form under the reaction conditions (Figure SI-30).

### Metho**ds**

#### General Methods

Reagents and solvents were obtained from *Acros Organics*, *Alfa Aesar*, *Sigma-Aldrich* and *VWR International*, and were used without further purification unless otherwise stated. For solid phase RNA synthesis, primer Support 5G for A, G, C, U (with loading ∼300 μmol/g) was purchased from *GE Healthcare*. Phosphoramidites for RNA synthesis were purchased from *Sigma-Aldrich* or *Link Technologies*. RNA oligomers used in this study were synthesized using an ÄKTA™ oligopilot™ plus 10 (*GE Healthcare*) on a 5 to 50 μmol scale. *MettlerToledo* SevenEasy pH Meter S20 combined with a *ThermoFisher Scientific* Orion 8103BN Ross semi-micro pH electrode was used to measure and adjust the pH to the desired value. ^1^H-, and ^31^P-nuclear magnetic resonance (NMR) spectra were acquired using a *Bruker* Ultrashield 400 Plus or *Bruker* Ascend 400 operating at 400.13, and 161.97 MHz, respectively. Samples consisting of H_2_O/D_2_O mixtures were analyzed using HOD suppression to collect ^1^H-NMR spectroscopy data. Chemical shifts (δ) are shown in ppm. Mass spectra were acquired on an *Agilent* 1200 LC-MS system equipped with an electrospray ionization (ESI) source and a 6130 quadrupole spectrometer (LC solvents: A, 0.2% formic acid in H_2_O – B, and 0.2% formic acid in acetonitrile), or on a *Bruker* Ultraflex III MALDI-TOF. High-Pressure Liquid Chromatography (HPLC) was run on Dionex Ultimate 3000 (*Thermo Scientific*) using an Atlantis^TM^ T3, 5 μm, 4.6 x 250 mm column or Atlantis^TM^ T3, 3 μm, 4.6 x 150 mm column. Oligonucleotide concentrations were determined by UV absorbance at 260 nm using a NanoDrop® ND-1000 spectrophotometer. Phosphoramidate-ester **4** and **7** yields were calculated by acid hydrolysis of **4** and **7**, and subsequent comparison against calibration curves of the products 3'-AGCGAp **6,** 3'-ACCUUUCGCU **3** and 3'-AAGGUAAU **8** (Charts SI-1 to SI-3). In most cases D- phosphoramidate-esters were isolated in extremely low yields, which made direct quantification impractical. In these cases we assumed the difference in UV absorbance between these D- and L- phosphoramidate-esters **4** or **7** was similar to that of other measurable D-/L- phosphoramidate-esters.

#### Chemical synthesis of RNA oligomers

After automated synthesis, RNAs were first cleaved from the solid support by treating with 3 mL of a 1:1 mixture of 28% wt NH_3_/H_2_O solution and 33% wt CH_3_NH_2_/EtOH solution at 65 °C for 90 minutes in a tube with a sealed cap. The solid was removed by filtration and washed with 50% EtOH/H_2_O. The solution and washings were combined and evaporated to remove all ethanol under reduced pressure. The residue was lyophilized to dryness. Silyl protecting groups were then removed by treating the residues with 3 mL of 1:1 mixture of triethylamine trihydrofluoride and DMSO at 65 °C for 180 minutes in a tube with a sealed cap. After brief cooling at -32 °C, 30 mL of cold 50 mM NaClO_4_ in acetone was added to the solution to precipitate the RNA product. The resulting mixture was centrifuged and the pellet of RNA was re-dissolved in 10 mL of water and passed through a *Waters* Sep-Pak C18 Cartridge, 10 g sorbent (Cartridge was pre-washed with 20 mL of MeOH then 100 mL of water before sample loading, then washed with 150 mL of H_2_O, 40 mL of 10% MeOH/H_2_O, 40 mL of 20% MeOH/H_2_O, 40 mL of 50% MeOH/H_2_O and 40 mL of MeOH sequentially). Eluents containing RNA were combined and if the tetrabutyl ammonium RNA salt was required, were neutralised with tetrabutyl ammonium hydroxide (~40% in water). The solutions were lyophilized and the resulting RNA was stored as a solid or dissolved in RNase-free water at -32 °C for future usage.

#### Chemical synthesis of RNA phosphoramidate (2): Method 1

4-(dimethylamino)pyridine (DMAP, 34.9 mg, 0.29 mmol) and triphenylphosphine (76.1 mg, 0.29 mmol) were dissolved in DMSO (250 uL). Solid tetrabutylammonium RNA pentamer **6** (3^'^AGCGAp, 4 mg, 2.4 μmol) was dissolved in the DMSO solution. Finally dipyridyl disulphide (63.9 mg, 0.29 mmol) was added and the solution was left at room temperature for 2 hours. The solution was added into 30 ml of cold 1:1 50 mM NaClO_4_ in acetone:diethyl ether to precipitate the 3^'^AGCGAp-DMAP. The precipitate was separated by centrifugation and washed with diethyl ether (30 ml x 2), then dried *in vacuo*. A solution of amino acid (50 mg/ml) was prepared and pH was adjusted to 8.0 by the addition of NaOH. From that, 0.5 ml solution was added to the dried precipitate and incubated at room temperature for 18 hours. The reaction was monitored by ^31^P-NMR spectroscope or HPLC. The products were purified by HPLC and stored as a solid or dissolved in basic pH solution at -32 °C for future usage. (Atlantis^TM^ T3, 3 μm, 4.6 x 150 mm column or Atlantis^TM^ T3, 5 μm, 4.6 x 250 mm column; flow rate 1 mL/min; flow rate 1 mL/min; LC solvents: A, 50 mM triethylammonium acetate, pH 7 in water and B, acetonitrile. Gradient: for L-Proline 0 min (7% B), to 1 min (7% B), to 9 min (8% B), and 9.3 min (85% B) or for L-leucine, L-valine and D-leucine 0 min (7% B), to 1 min (7% B), to 15 min (14% B) and 15.3 min (85% B) or for L-serine, D-serine, L-arginine and D-alanine 0 min (7% B), to 1 min (7% B), to 9 min (12% B) and 9.2 min (85% B) or for D-valine and L-leucine 0 min (7% B), to 1 min (7% B), to 15 min (14% B), and 15.3 min (85% B) or for L-alanine 0 min (7% B), to 1 min (7% B), to 20 min (12% B), to 22 min (16% B) and 23 min (95% B). Column compartment temperature at 25 °C).

#### Chemical synthesis of RNA phosphoramidate (2)^12^: Method 2

4-(dimethylamino)pyridine (DMAP, 6.6 mg, 0.054 mmol) and RNA pentamer **6** (3^'^AGCGAp, 3 mg, 1.8 μmol) were dissolved in H_2_O/D_2_O (9:1, *v*/*v*, 0.5 ml), the pH value was adjusted to 8.0 with hydrochloric acid (5 M). To the resulting solution, 1-ethyl-3-(3-dimethylaminopropyl)carbodiimide hydrochloride (EDC, 50 mg, 0.23 mmol) was added. The reaction mixture was kept at room temperature for 2 hours, and the reaction was followed by ^31^P-NMR spectroscope. Afterwards, the solution was added dropwise into 10 ml of cold 50 mM NaClO_4_ in acetone to precipitate the 3^'^AGCGAp-DMAP. The precipitate was separated by centrifugation and washed with diethyl ether (10 ml), then dried *in vacuo*. 3^'^AGCGAp-DMAP was then converted into phosphoramidates **2** as Method 1.

#### The formation of RNA phosphoramidate-esters (4) or (7)

A mixture containing the above synthesized RNA phosphoramidate **2** with amino acid (100 μM,), aminoacyl acceptor RNA **3** (100 μM, ^3'^ACCUUUCGCU), complimentary RNA **8** (if required, 100 μM, ^3'^AAGGUAAU), NaCl (200 mM), MgCl_2_ (50 mM), EDC (50 mM) and imidazole (10 mM) in HEPES buffer (100 mM, pH 7) with cytidine (120 μM) as an internal standard if required, was incubated at the desired temperature and time. Aliquots (2 μl) were taken at specific time points and diluted in RNase-free water. The resultant mixture (5 μl) was injected directly to an HPLC for analysis at 260 nm UV detection (Atlantis^TM^ T3, 5 μm, 4.6 x 250 mm column; flow rate 1 mL/min; LC solvents: A, 20 mM triethylammonium acetate, pH 7 in water and B, acetonitrile. Gradient: 0 min (7% B), to 1 min (7% B), to 20 min (12% B), to 22 min (16% B) and 23 min (95% B). Column compartment temperature at 25 °C).

#### Phosphoramidate-ester (4) purification

RNA phosphoramidate-ester **4** was purified by HPLC (Atlantis^TM^ T3, 3 μm, 4.6 x 150 mm column; flow rate 1 mL/min; LC solvents: A, 20 mM triethylammonium acetate, pH 7 in water and B, acetonitrile. Gradient: 0 min (7% B), to 1 min (7% B), to 9 min (8% B), and 9.3 min (85% B) or 0 min (7% B), to 1 min (7% B), to 13 min (12% B), to 14 min (16%) and 14.2 min (95% B) or 0 min (7% B), to 1 min (7% B), to 9 min (12% B), to 10 min (16%) and 10.2 min (95% B). Column compartment temperature at 25 °C. The resultant solution was dried by lyophilization.

#### **Acid hydrolysis of phosphoramidate-ester (4**)

The lyophilised RNA phosphoramidate-ester **4** was dissolved in water (10 μL). An aliquot of phosphoramidate solution (5 μL) was combined with formate buffer (25 μL, 100 mM, pH 3) containing cytidine as internal standard (80 μM) and incubated at 25 °C for 17 hours. Samples were then directly analysed by HPLC for analysis at 260 nm UV detection (Atlantis^TM^ T3, 5 μm, 4.6 x 250 mm column; flow rate 1 mL/min; LC solvents: A, 20 mM triethylammonium acetate, pH 7 in water and B, acetonitrile. Gradient: 0 min (7% B), to 1 min (7% B), to 20 min (12% B), to 22 min (16% B) and 23 min (95% B). Column compartment temperature at 25 °C) and then incubated at 25 °C for 17 h before further analysis by HPLC (method as above).

#### **Base hydrolysis of phosphoramidate-ester (4)**^15^

NaOH (100 mM, 15 μl) solution was added to the crude RNA phosphoramidate-ester **4** formation solution (15 μl). After incubation for 60 s, HCl (105 mM, 15 μl) solution was added to the resultant solution altering the pH value of the solution to around 5.0. The sample was injected directly to an HPLC for analysis at 260 nm UV detection (Atlantis^TM^ T3, 5 μm, 4.6 x 250 mm column; flow rate 1 mL/min; LC solvents: A, 20 mM triethylammonium acetate, pH 7 in water and B, acetonitrile. Gradient: 0 min (7% B), to 1 min (7% B), to 20 min (12% B), to 22 min (16% B) and 23 min (95% B). Column compartment temperature at 25 °C).

#### Base hydrolysis of acid treated phosphoramidate-ester (4)

NaOH (1 M, 3 μl) solution was added to the crude formate treated RNA phosphoramidate-ester **4** (20 μl). After incubation for at least 60 s, HCl (2 M, 1.5 μl) solution was added to the result solution. The sample was injected directly to an HPLC for analysis at 260 nm UV detection (Atlantis^TM^ T3, 5 μm, 4.6 x 250 mm column; flow rate 1 mL/min; LC solvents: A, 20 mM triethylammonium acetate, pH 7 in water and B, acetonitrile. Gradient: 0 min (7% B), to 1 min (7% B), to 20 min (12% B), to 22 min (16% B) and 23 min (95% B). Column compartment temperature at 25 °C).

### Experimental Data

#### Phosphoramidate-ester (4) formation

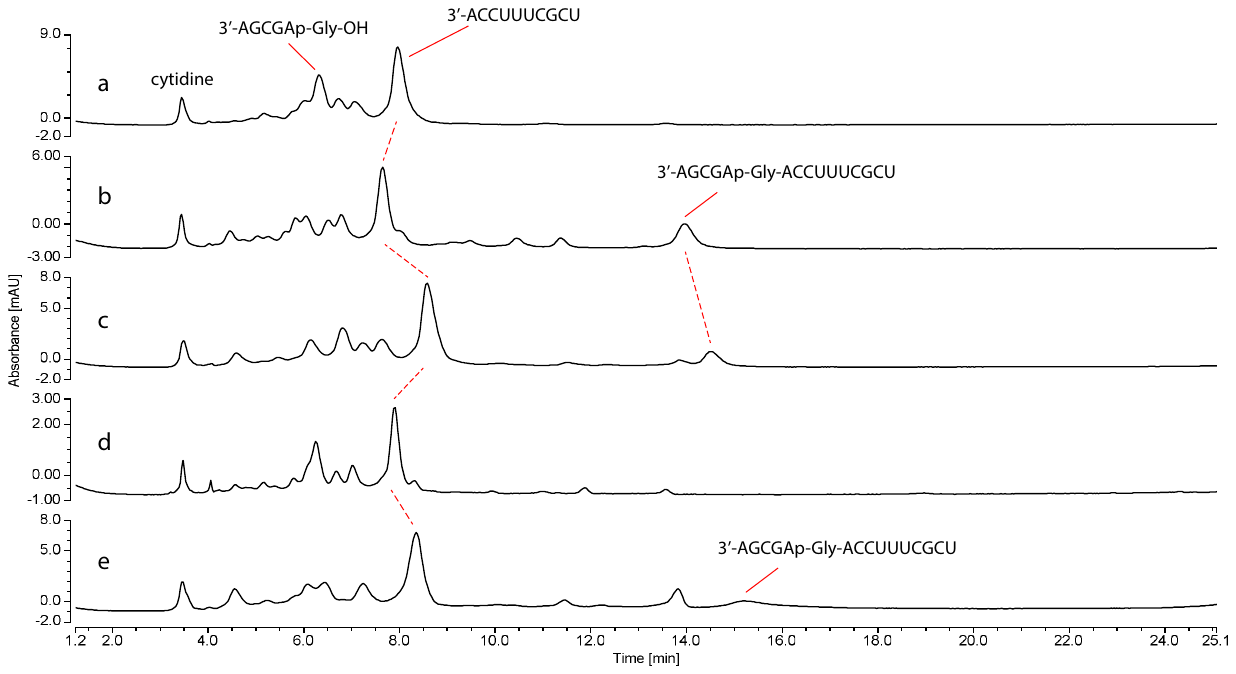

##### Figure SI-1. HPLC traces of the formation of RNA-glycine phosphoramidate-ester (4-Gly).

Loop duplex sequence:

3’AGCGAp-Gly-OH

5’UCGCUUUCCA

Reactions were monitored using HPLC with 260 nm UV detection. The solution was divided into aliquots which were either incubated at 20 °C for 18 hours or at -16 °C for 7 or 14 days. After the desired time each aliquot was diluted in 18 μL water, the diluted solutions were injected into an HPLC. a. Reaction after 0 hours; b. Reaction after 18 hours at room temperature; c. Reaction after 7 days at -16 °C; d. Sample c after base hydrolysis; e. Reaction after 14 days at -16 °C.

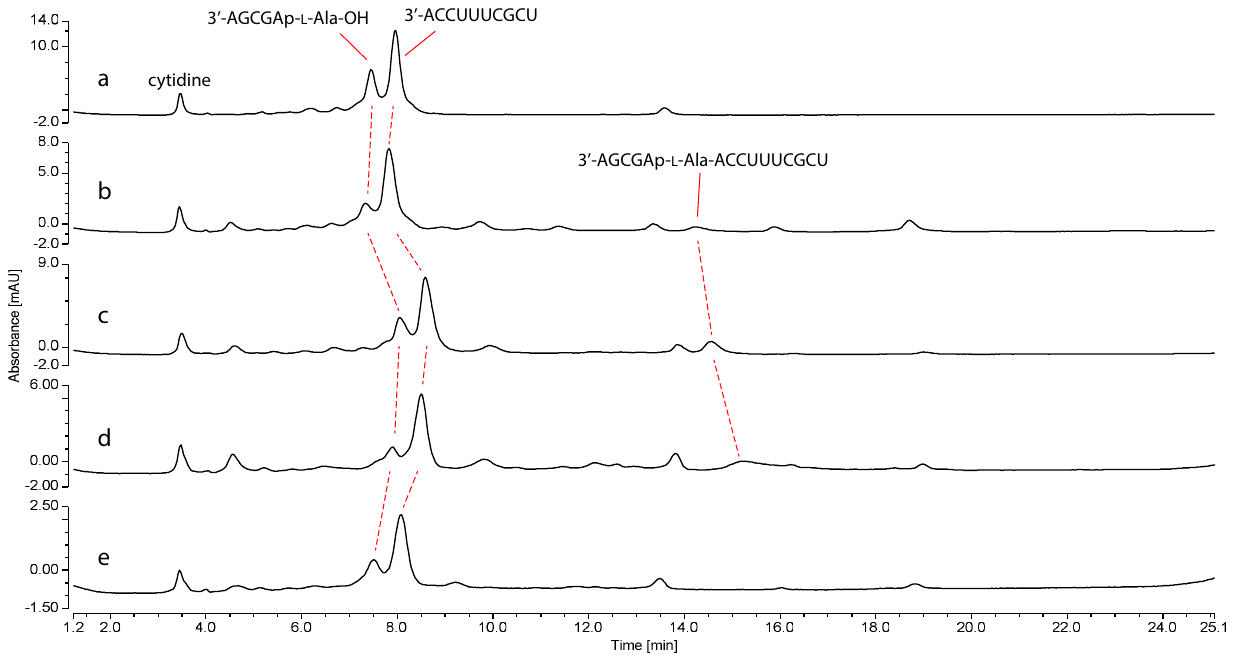

##### Figure SI-2. HPLC traces of the formation of RNA-L-Alanine phosphoramidate-ester (4- L -Ala).

Loop duplex sequence:

3’AGCGAp-L-Ala-OH

5’UCGCUUUCCA

Reactions were monitored using HPLC with 260 nm UV detection. The solution was divided into aliquots which were either incubated at 20 °C for 18 hours or at -16 °C for 7 or 14 days. After the desired time each aliquot was diluted in 18 μL water, the diluted solutions were injected into an HPLC. a. Reaction after 0 hours; b. Reaction after 18 hours at room temperature; c. Reaction after 7 days at -16 °C; d. Reaction after 14 days at -16 °C; e. Sample d after base hydrolysis.

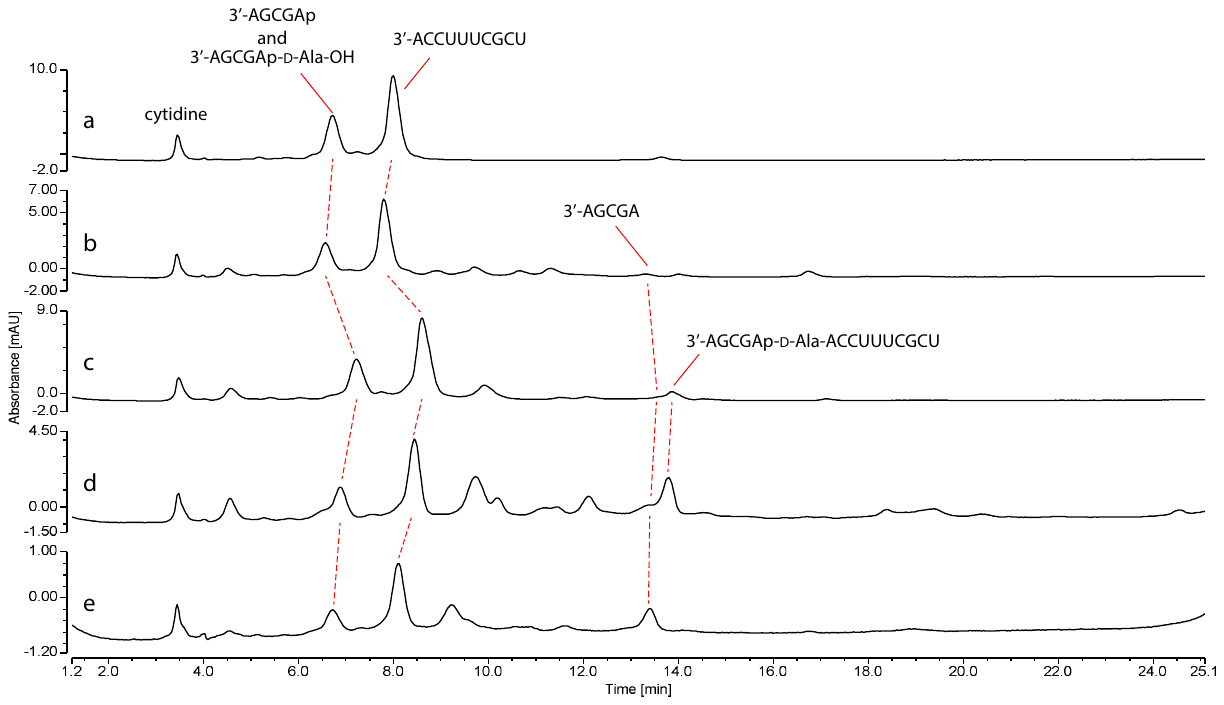

##### Figure SI-3. HPLC traces of the formation of RNA-D-Alanine phosphoramidate-ester (4-D-Ala).

Loop duplex sequence:

3’AGCGAp-D-Ala-OH

5’UCGCUUUCCA

Reactions were monitored using HPLC with 260 nm UV detection. The solution was divided into aliquots which were either incubated at 20 °C for 18 hours or at -16 °C for 7 or 14 days. After the desired time each aliquot was diluted in 18 μL water, the diluted solutions were injected into an HPLC. a. Reaction after 0 hours; b. Reaction after 18 hours at room temperature; c. Reaction after 7 days at -16 °C; d. Reaction after 14 days at -16 °C; e. Sample d after base hydrolysis.

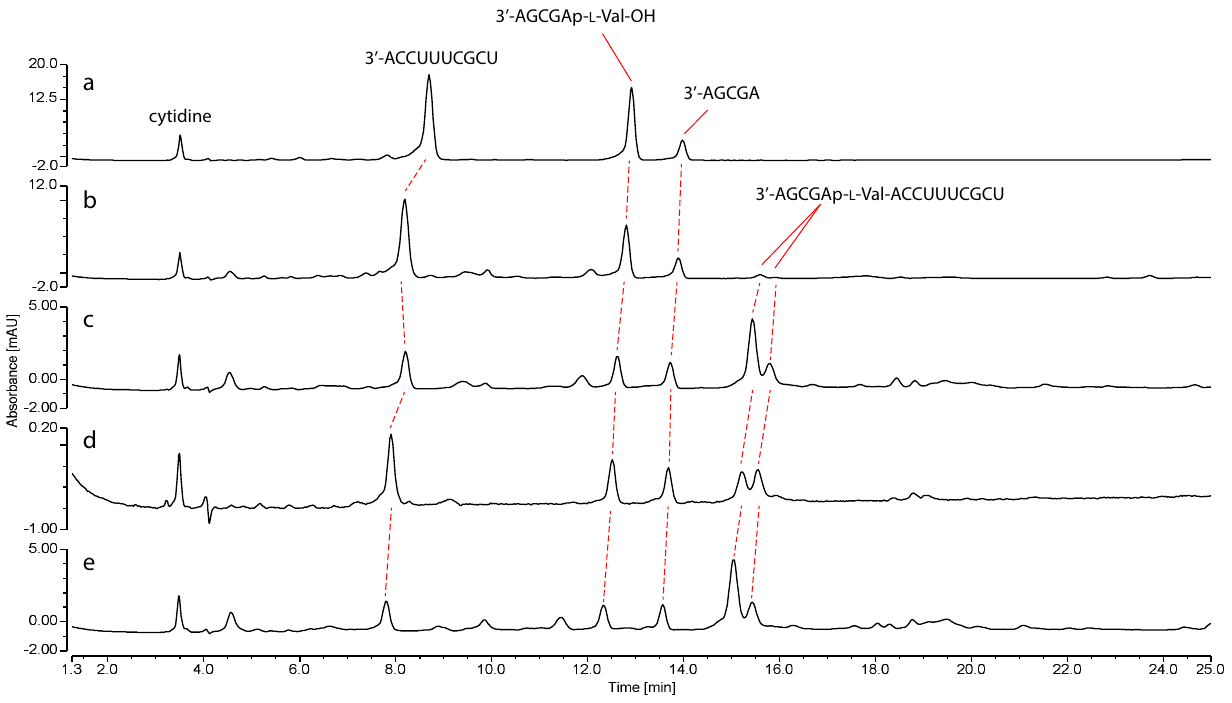

##### Figure SI-4. HPLC traces of the formation of RNA-L-Valine phosphoramidate-ester (4-L-Val).

Loop duplex sequence:

3’AGCGAp-L-Val-OH

5’UCGCUUUCCA

Reactions were monitored using HPLC with 260 nm UV detection. The solution was divided into aliquots which were either incubated at 20 °C for 18 hours or at -16 °C for 7 or 14 days. After the desired time each aliquot was diluted in 18 μL water, the diluted solutions were injected into an HPLC. a. Reaction after 0 hours; b. Reaction after 18 hours at room temperature; c. Reaction after 7 days at -16 °C; d. Sample c after base hydrolysis; e. Reaction after 14 days at -16 °C.

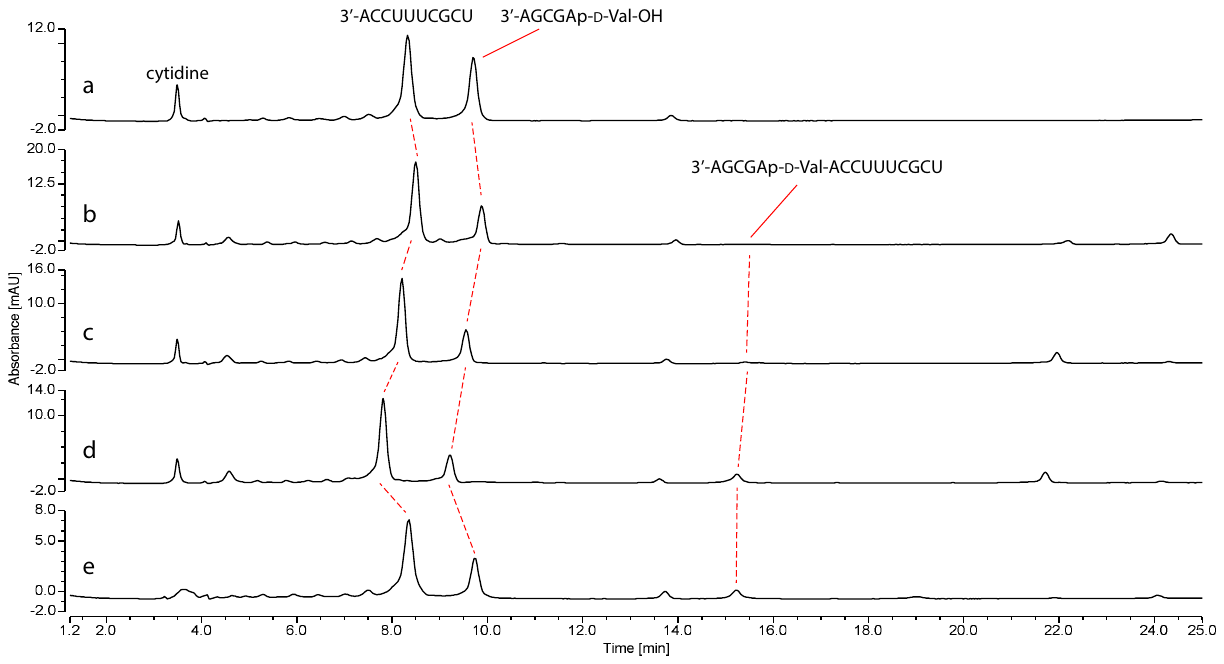

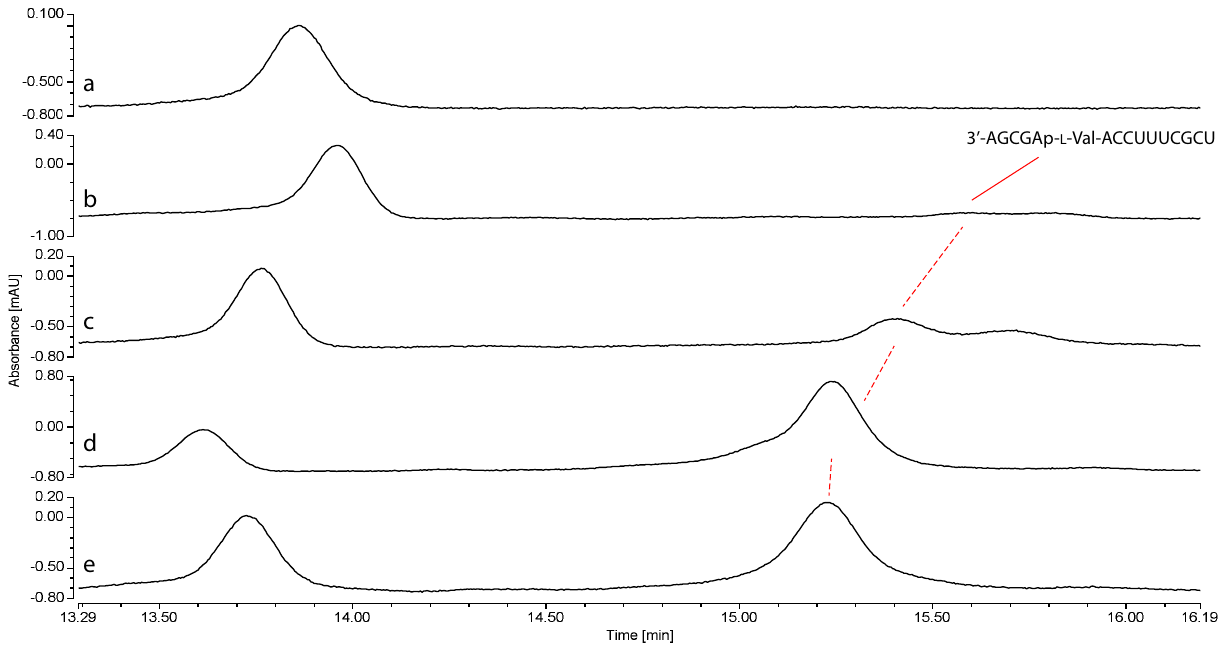

##### Figure SI-5. HPLC traces of the formation of RNA-D-Valine phosphoramidate-ester (4-D-Val). Top – Full chromatogram. Bottom – Zoom in around phosphoramidate-ester (4-D-Val) peak.

Loop duplex sequence:

3’AGCGAp-D-Val-OH

5’UCGCUUUCCA

Reactions were monitored using HPLC with 260 nm UV detection. The solution was divided into aliquots which were either incubated at 20 °C for 18 hours or at -16 °C for 7 or 14 days. After the desired time each aliquot was diluted in 18 μL water, the diluted solutions were injected into an HPLC. a. Reaction after 0 hours; b. Reaction after 18 hours at room temperature; c. Reaction after 7 days at -16 °C; d. Reaction after 14 days at -16 °C; e. Sample d after base hydrolysis.

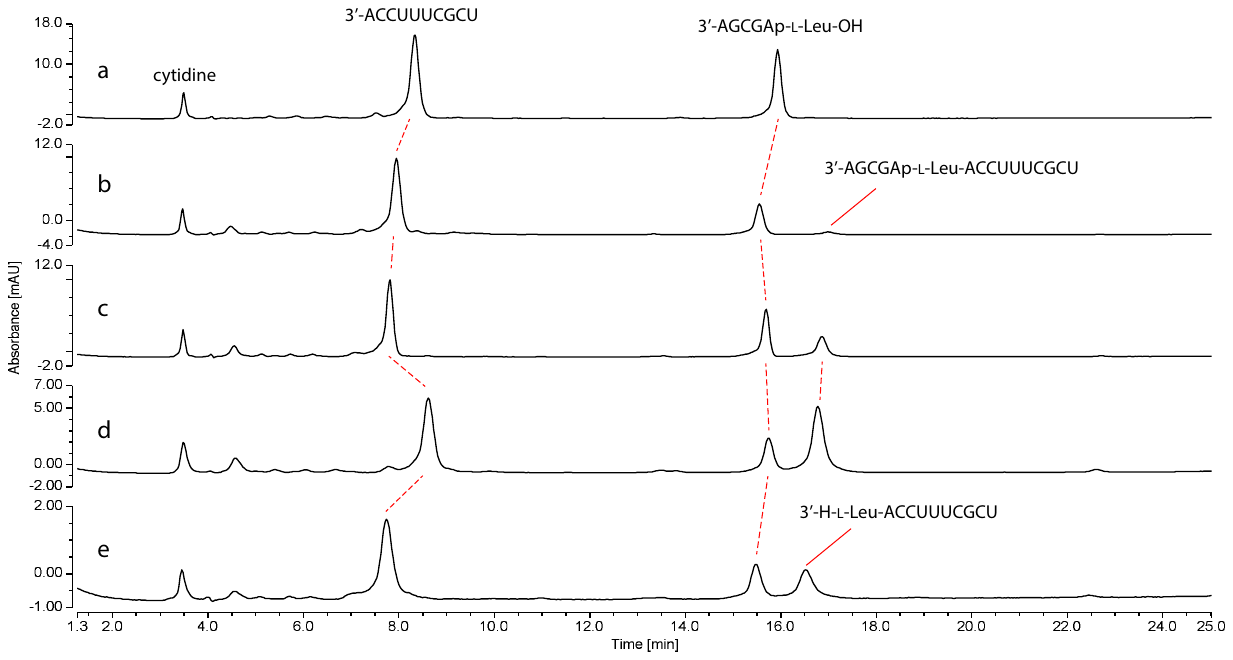

##### Figure SI-6. HPLC traces of the formation of RNA-L-Leucine phosphoramidate-ester (4-L-Leu).

Loop duplex sequence:

3’AGCGAp-L-Leu-OH

5’UCGCUUUCCA

Reactions were monitored using HPLC with 260 nm UV detection. The solution was divided into aliquots which were either incubated at 20 °C for 18 hours or at -16 °C for 7 or 14 days. After the desired time each aliquot was diluted in 18 μL water, the diluted solutions were injected into an HPLC. a. Reaction after 0 hours; b. Reaction after 18 hours at room temperature; c. Reaction after 7 days at -16 °C; d. Reaction after 14 days at -16 °C; e. Sample d after base hydrolysis.

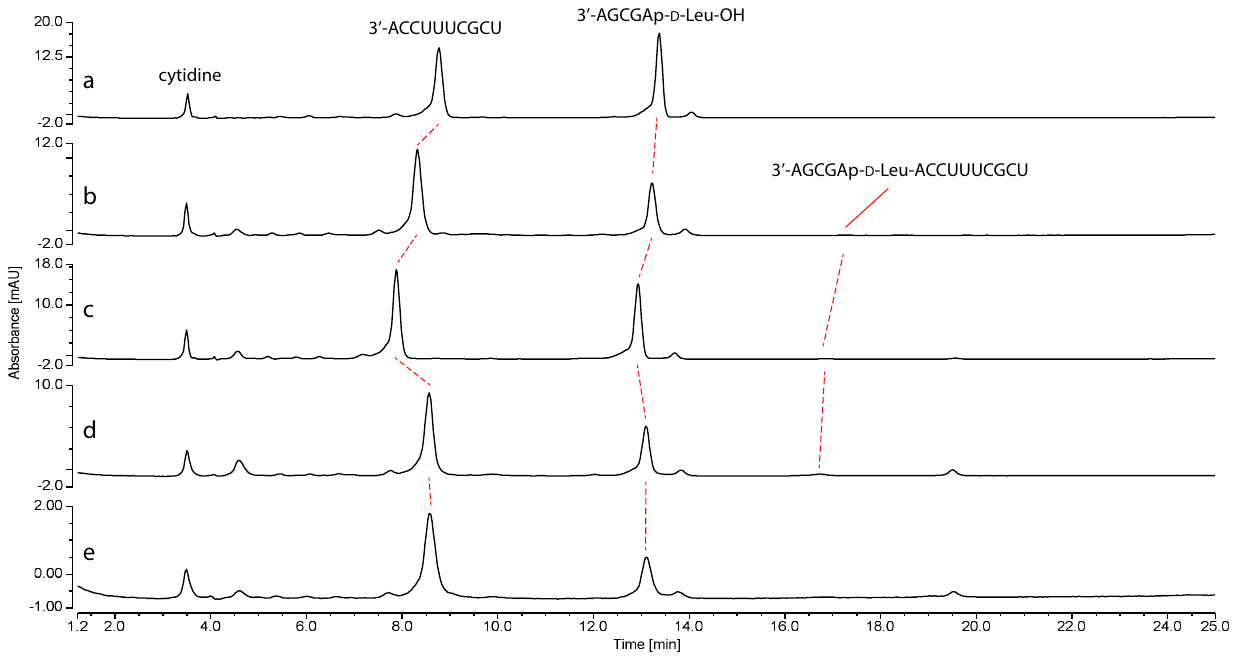

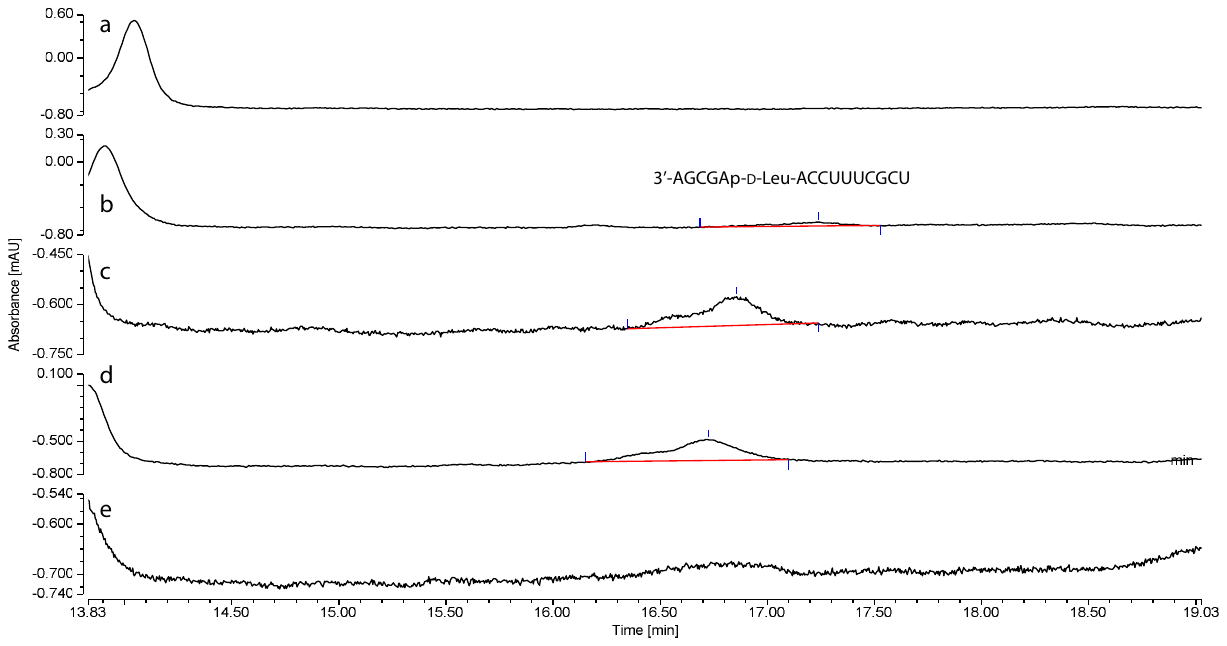

##### Figure SI-7. HPLC traces of the formation of RNA-D-Leucine phosphoramidate-ester (4-D-Leu). Top – Full chromatogram. Bottom – Zoom in around phosphoramidate-ester (4-D-Leu) peak.

Loop duplex sequence:

3’AGCGAp-D-Leu-OH

5’UCGCUUUCCA

Reactions were monitored using HPLC with 260 nm UV detection. The solution was divided into aliquots which were either incubated at 20 °C for 18 hours or at -16 °C for 7 or 14 days. After the desired time each aliquot was diluted in 18 μL water, the diluted solutions were injected into an HPLC. a. Reaction after 0 hours; b. Reaction after 18 hours at room temperature; c. Reaction after 7 days at -16 °C; d. Reaction after 14 days at -16 °C; e. Sample d after base hydrolysis.

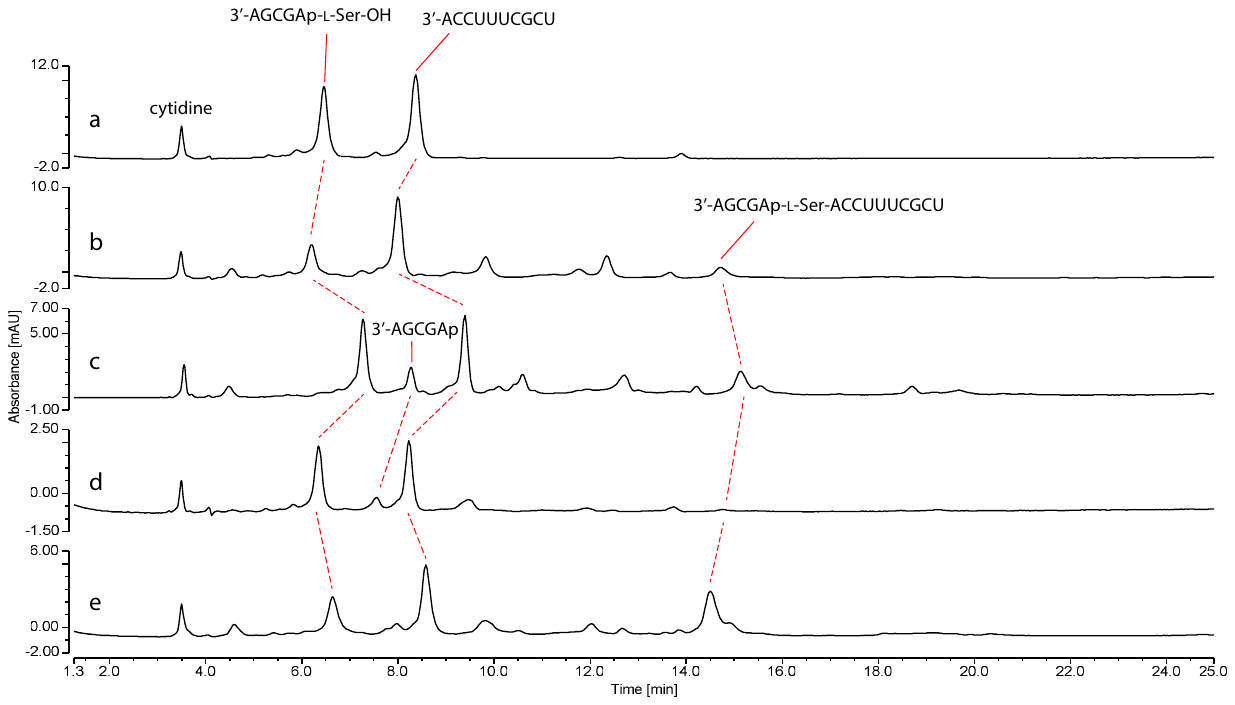

##### Figure SI-8. HPLC traces of the formation of RNA-L-Serine phosphoramidate-ester (4-L-Ser).

Loop duplex sequence:

3’AGCGAp-L-Ser-OH

5’UCGCUUUCCA

Reactions were monitored using HPLC with 260 nm UV detection. The solution was divided into aliquots which were either incubated at 20 °C for 18 hours or at -16 °C for 7 or 14 days. After the desired time each aliquot was diluted in 18 μL water, the diluted solutions were injected into an HPLC. a. Reaction after 0 hours; b. Reaction after 18 hours at room temperature; c. Reaction after 7 days at -16 °C; d. Sample c after base hydrolysis; e. Reaction after 14 days at -16 °C.

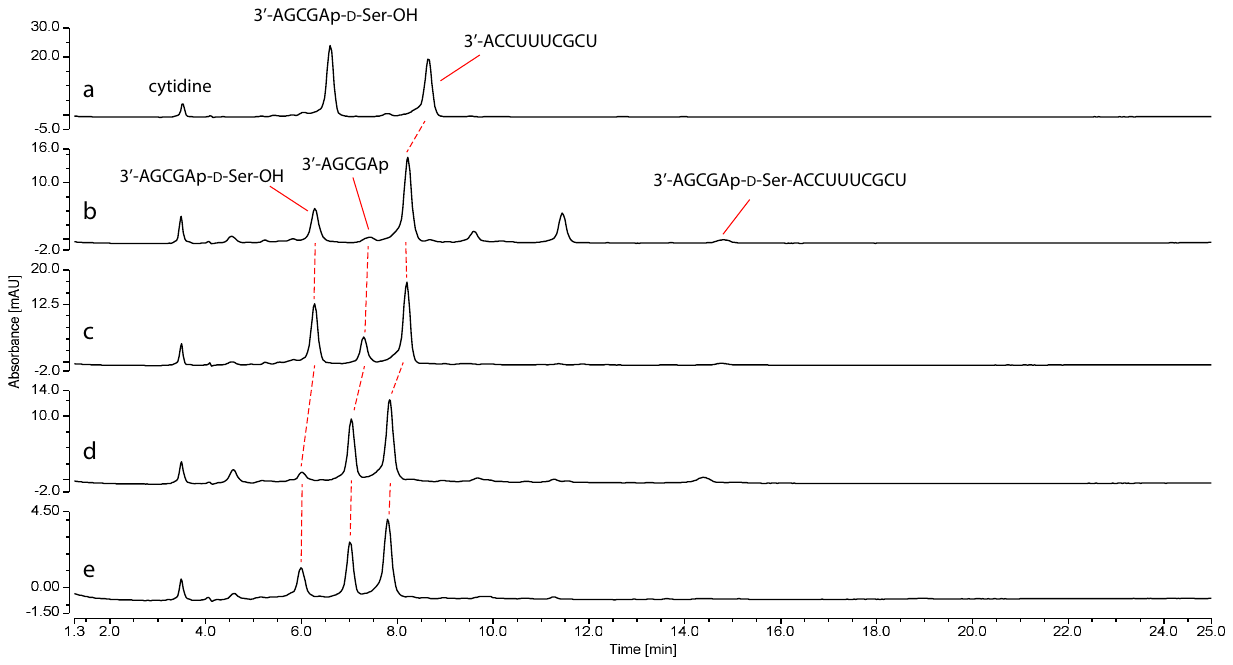

##### Figure SI-9. HPLC traces of the formation of RNA-D-Serine phosphoramidate-ester (4-D-Ser).

Loop duplex sequence:

3’AGCGAp-D-Ser-OH

5’UCGCUUUCCA

Reactions were monitored using HPLC with 260 nm UV detection. The solution was divided into aliquots which were either incubated at 20 °C for 18 hours or at -16 °C for 7 or 14 days. After the desired time each aliquot was diluted in 18 μL water, the diluted solutions were injected into an HPLC. a. Reaction after 0 hours; b. Reaction after 18 hours at room temperature; c. Reaction after 7 days at -16 °C; d. Reaction after 14 days at -16 °C; e. Sample d after base hydrolysis.

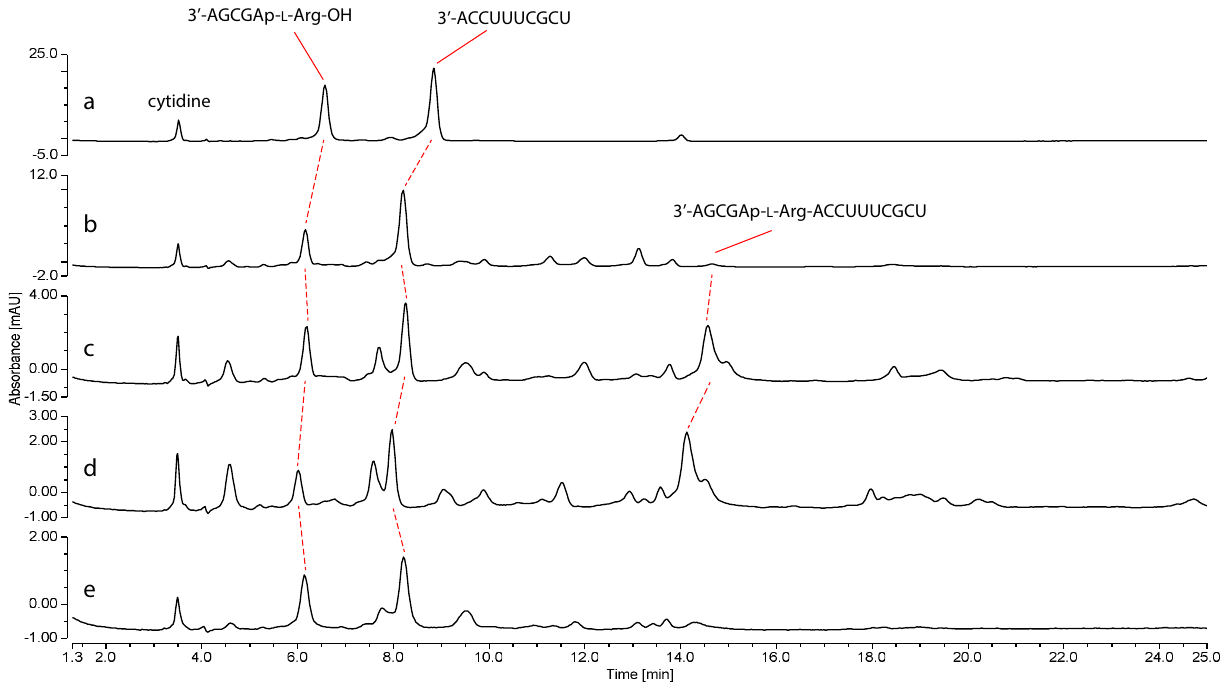

##### Figure SI-10. HPLC traces of the formation of RNA-L-Arginine phosphoramidate-ester (4-L-Arg).

Loop duplex sequence:

3’AGCGAp-L-Arg-OH

5’UCGCUUUCCA

Reactions were monitored using HPLC with 260 nm UV detection. The solution was divided into aliquots which were either incubated at 20 °C for 18 hours or at -16 °C for 7 or 14 days. After the desired time each aliquot was diluted in 18 μL water, the diluted solutions were injected into an HPLC. a. Reaction after 0 hours; b. Reaction after 18 hours at room temperature; c. Reaction after 7 days at -16 °C; d. Reaction after 14 days at -16 °C; e. Sample d after base hydrolysis.

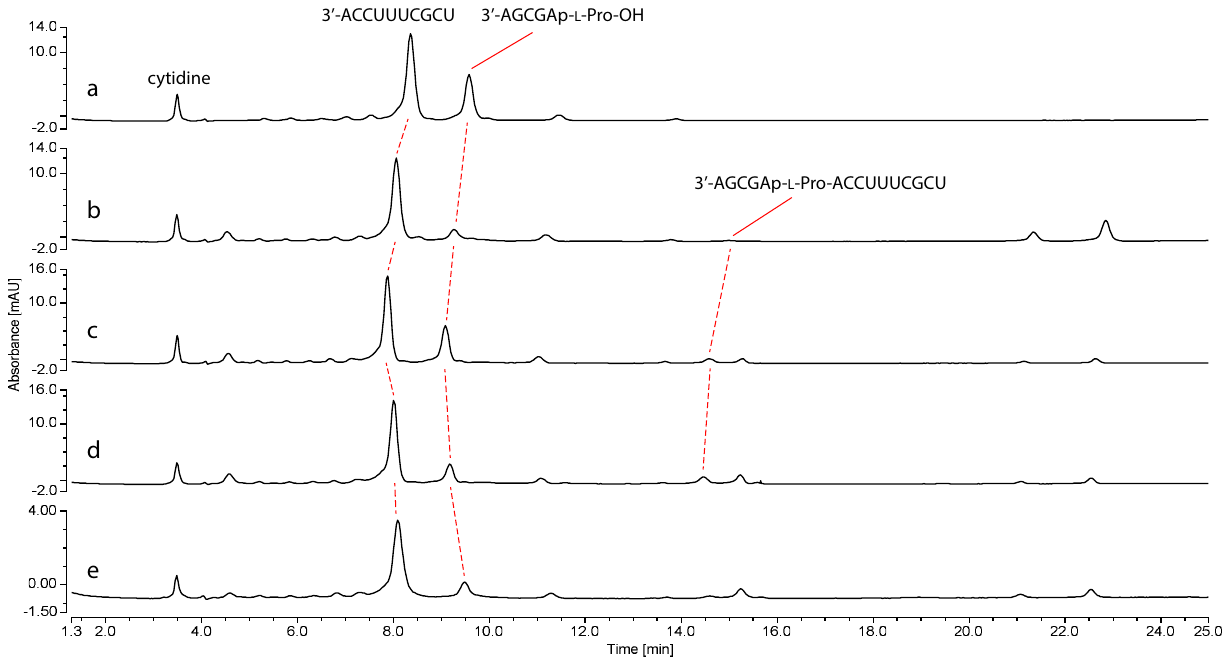

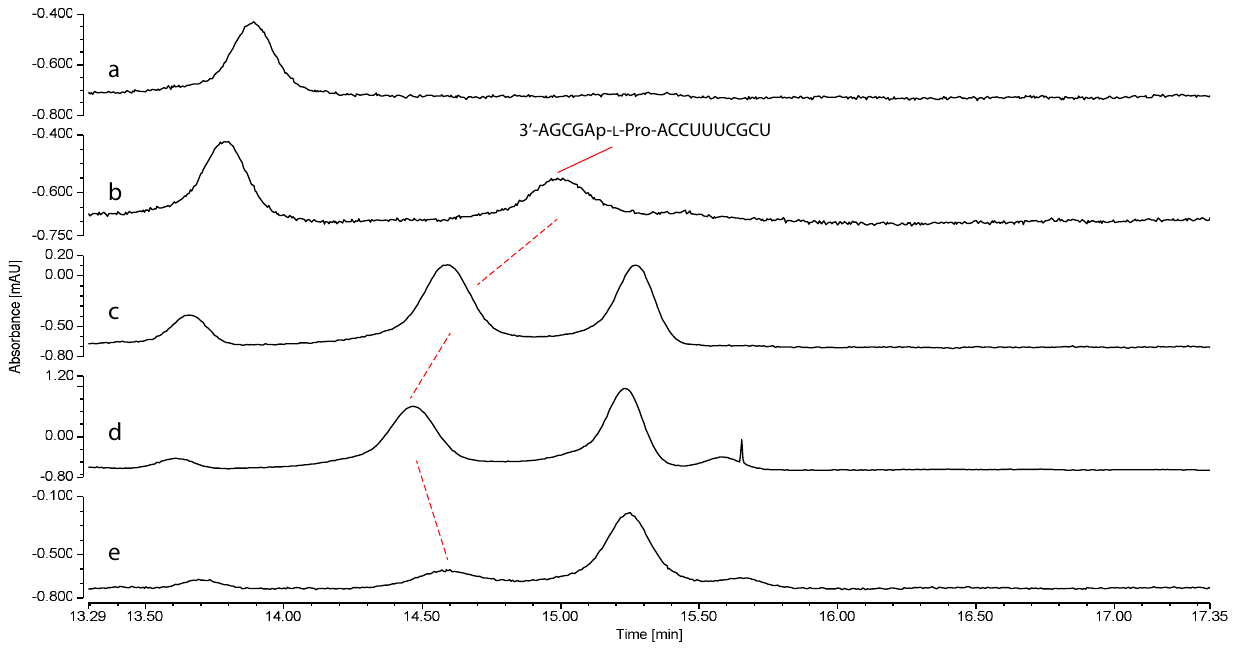

##### Figure SI-11. HPLC traces of the formation of RNA-L-Proline phosphoramidate-ester (4-L-Pro). Top – Full chromatogram. Bottom – Zoom in around phosphoramidate-ester (4-L-Pro) peak.

Loop duplex sequence:

3’AGCGAp-L-Pro-OH

5’UCGCUUUCCA

Reactions were monitored using HPLC with 260 nm UV detection. The solution was divided into aliquots which were either incubated at 20 °C for 18 hours or at -16 °C for 7 or 14 days. After the desired time each aliquot was diluted in 18 μL water, the diluted solutions were injected into an HPLC. a. Reaction after 0 hours; b. Reaction after 18 hours at room temperature; c. Reaction after 7 days at -16 °C; d. Reaction after 14 days at -16 °C; e. Sample d after base hydrolysis.

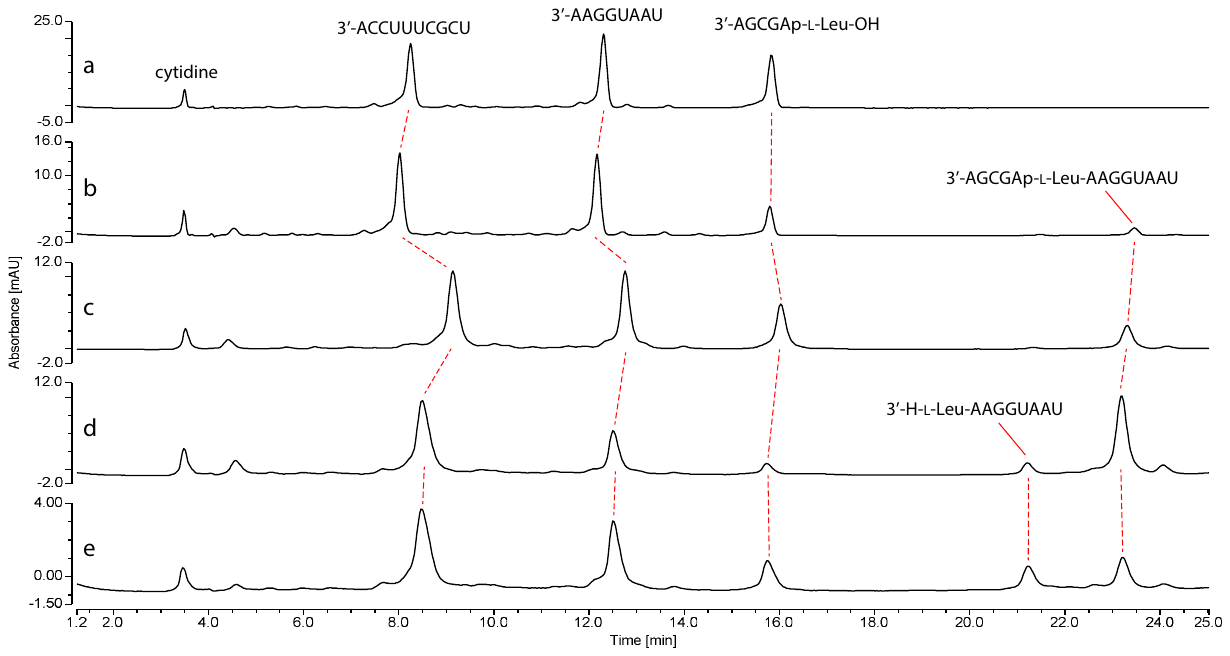

##### Figure SI-12. HPLC traces of the formation of nicked duplex RNA-L-Leucine phosphoramidate-ester (7-L-Leu).

Nicked duplex sequence:

3’AGCGAp-L-Leu-OH

5’UCGCUUUCCA

3’AAGGUAAU

Reactions were monitored using HPLC with 260 nm UV detection. The solution was divided into aliquots which were either incubated at 20 °C for 18 hours or at -16 °C for 7 or 14 days. After the desired time each aliquot was diluted in 18 μL water, the diluted solutions were injected into an HPLC. a. Reaction after 0 hours; b. Reaction after 18 hours at room temperature; c. Reaction after 7 days at -16 °C; d. Reaction after 14 days at -16 °C; e. Sample d after base hydrolysis.

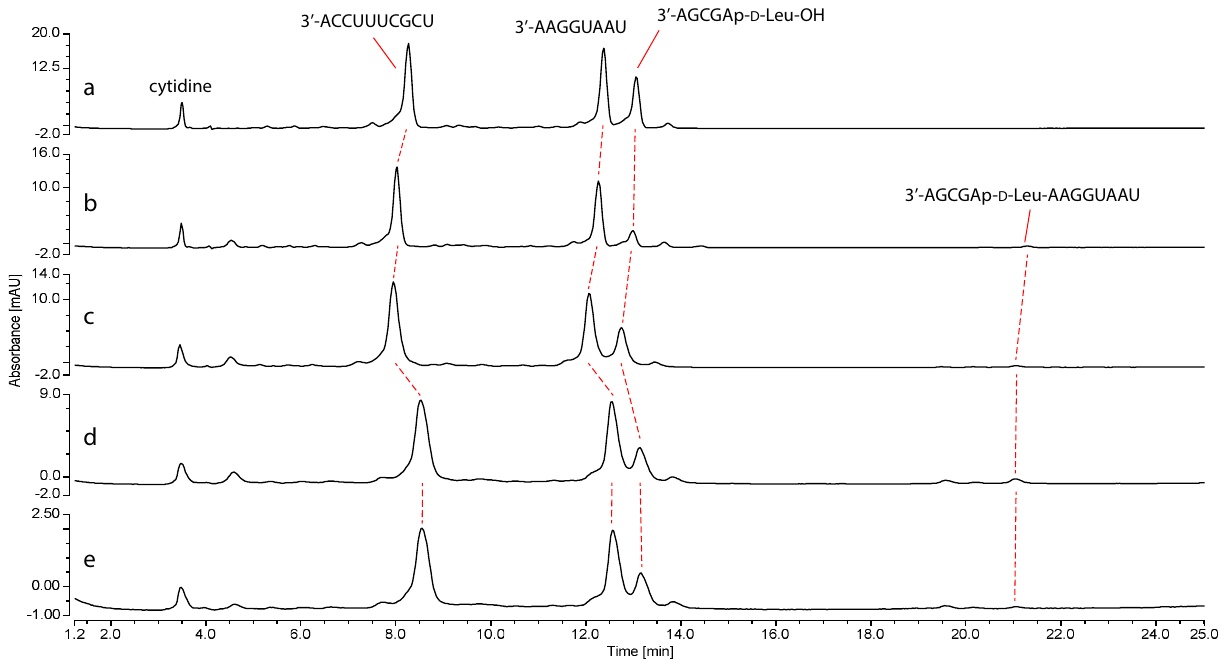

##### Figure SI-13. HPLC traces of the formation of nicked duplex RNA-D-Leucine phosphoramidate-ester (7-D-Leu).

Nicked duplex sequence:

3’AGCGAp-D-Leu-OH

5’UCGCUUUCCA

3’AAGGUAAU

Reactions were monitored using HPLC with 260 nm UV detection. The solution was divided into aliquots which were either incubated at 20 °C for 18 hours or at -16 °C for 7 or 14 days. After the desired time each aliquot was diluted in 18 μL water, the diluted solutions were injected into an HPLC. a. Reaction after 0 hours; b. Reaction after 18 hours at room temperature; c. Reaction after 7 days at -16 °C; d. Reaction after 14 days at -16 °C; e. Sample d after base hydrolysis.

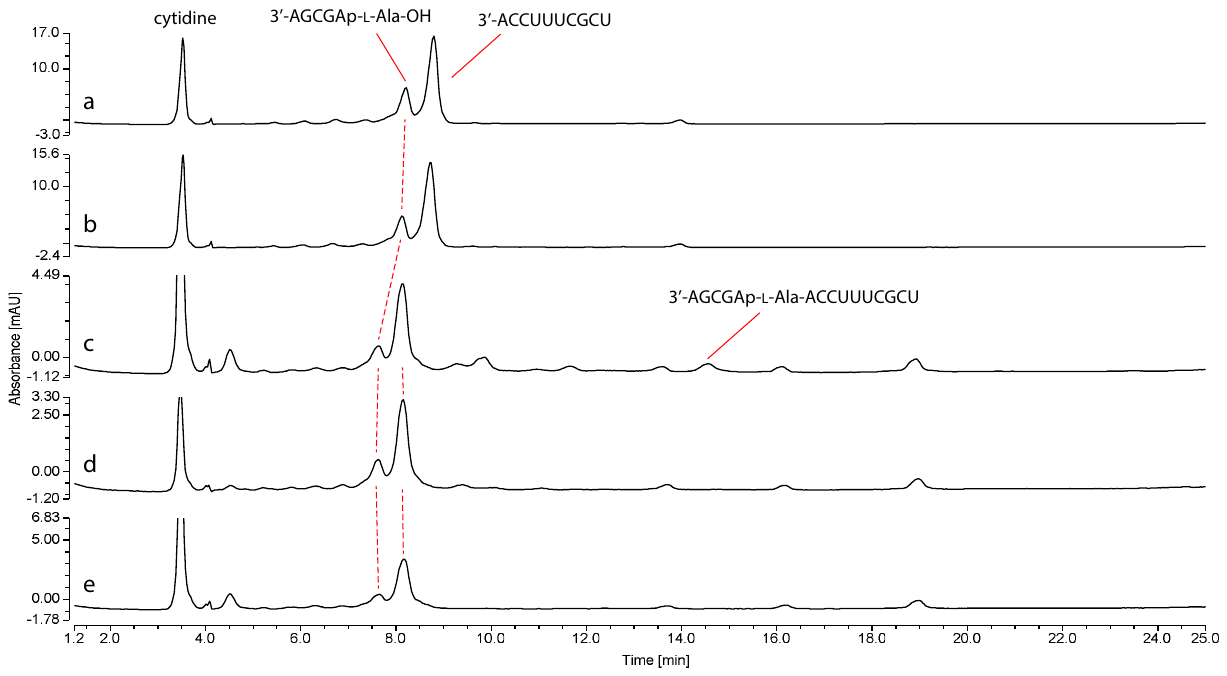

##### Figure SI-14. HPLC traces of the formation (or lack of formation) of RNA-L-Alanine phosphoramidate-ester (4-L-Ala) in the presence/absence of imidazole.

Loop duplex sequence:

3’AGCGAp-L-Ala-OH

5’UCGCUUUCCA

Reactions were monitored using HPLC with 260 nm UV detection. The solution was divided into aliquots which were either incubated at 20 °C for 18 hours or at -16 °C for 7 or 14 days. After the desired time each aliquot was diluted in 18 μL water, the diluted solutions were injected into an HPLC. a. Reaction after 0 hours with imidazole; b. Reaction after 0 hours without imidazole; c. Reaction after 18 hours at room temperature with imidazole; d. Sample c after base hydrolysis; e. Reaction after 18 hours at room temperature without imidazole.

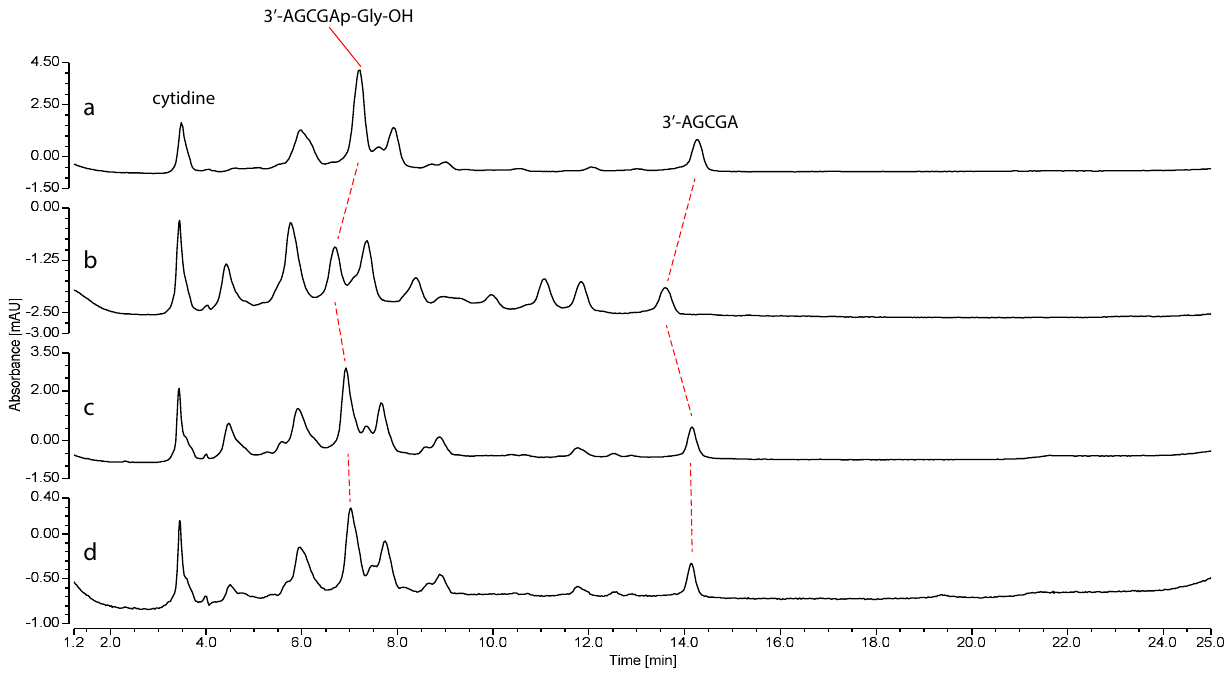

##### Figure SI-15. HPLC traces of the control reaction of Glycine amidate (2-Gly).

RNA sequence:

3’AGCGAp-Gly-OH

Reactions were monitored using HPLC with 260 nm UV detection. The solution was divided into aliquots which were either incubated at 20 °C for 18 hours or at -16 °C for 7 or 14 days. After the desired time each aliquot was diluted in 18 μL water, the diluted solutions were injected into an HPLC. a. Reaction after 0 hours; b. Reaction after 18 hours at room temperature; c. Reaction after 7 days at -16 °C; d. Sample c after base hydrolysis.

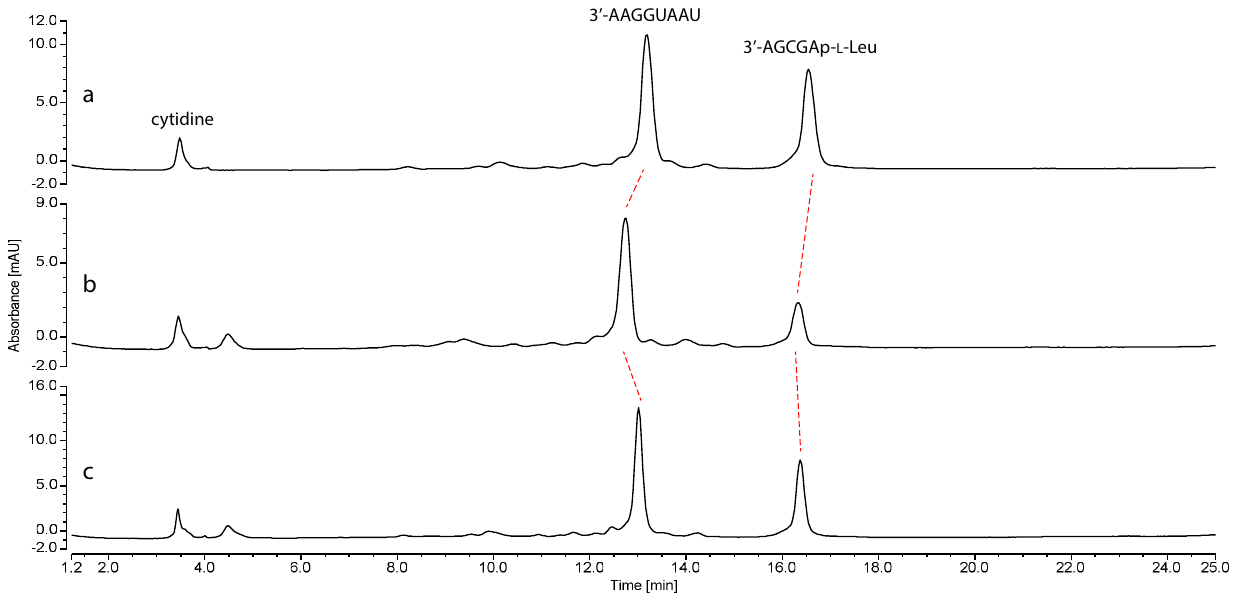

##### Figure SI-16. HPLC traces of the control reaction of Leucine amidate (2-L-Leu) with 8mer (8).

RNA sequence:

3’AGCGAp-L-Leu-OH

5’UAAUGGAA

Reactions were monitored using HPLC with 260 nm UV detection. The solution was incubated under the desired conditions and aliquots of 2 μL of the reaction solution were diluted in 18 μL water, the diluted solutions were injected into an HPLC. a. Reaction after 0 hours; b. Reaction after 18 hours at room temperature; c. Reaction after 7 days at -16 °C.

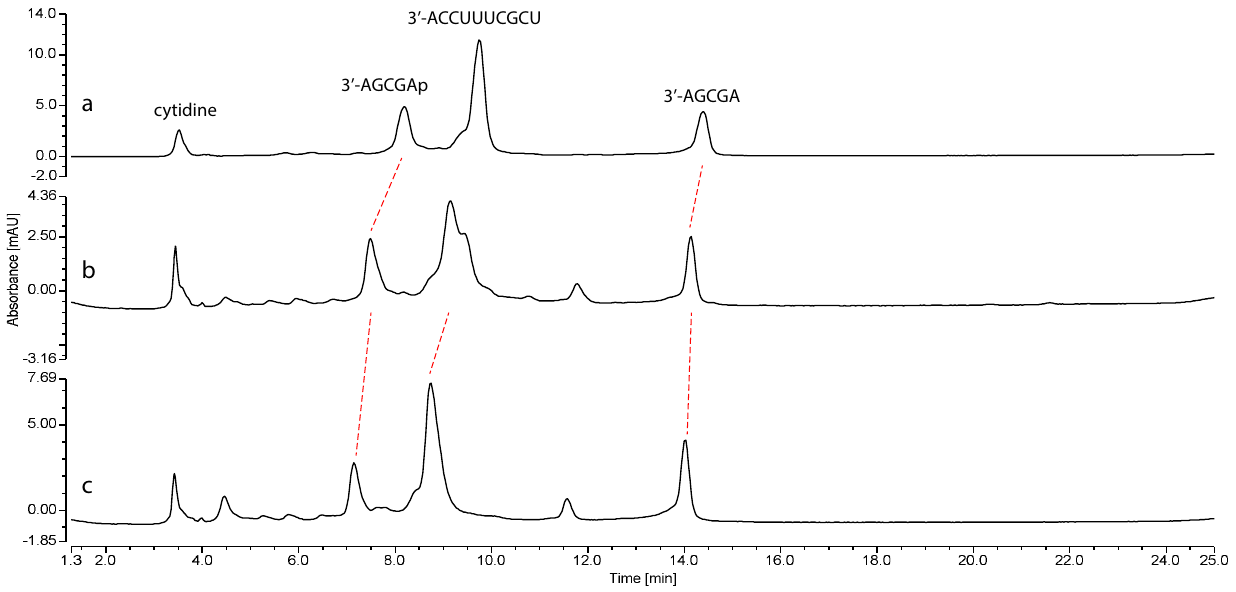

##### Figure SI-17. HPLC traces of the control reaction of 5'P-5mer (6) with 10mer (3).

Loop duplex sequence:

3’AGCGAp

5’UCGCUUUCCA

Reactions were monitored using HPLC with 260 nm UV detection. The solution was incubated under the desired conditions and aliquots of 2 μL of the reaction solution were diluted in 18 μL water, the diluted solutions were injected into an HPLC. a. Reaction after 0 hours; b. Reaction after 18 hours at room temperature; c. Reaction after 7 days at -16 °C.

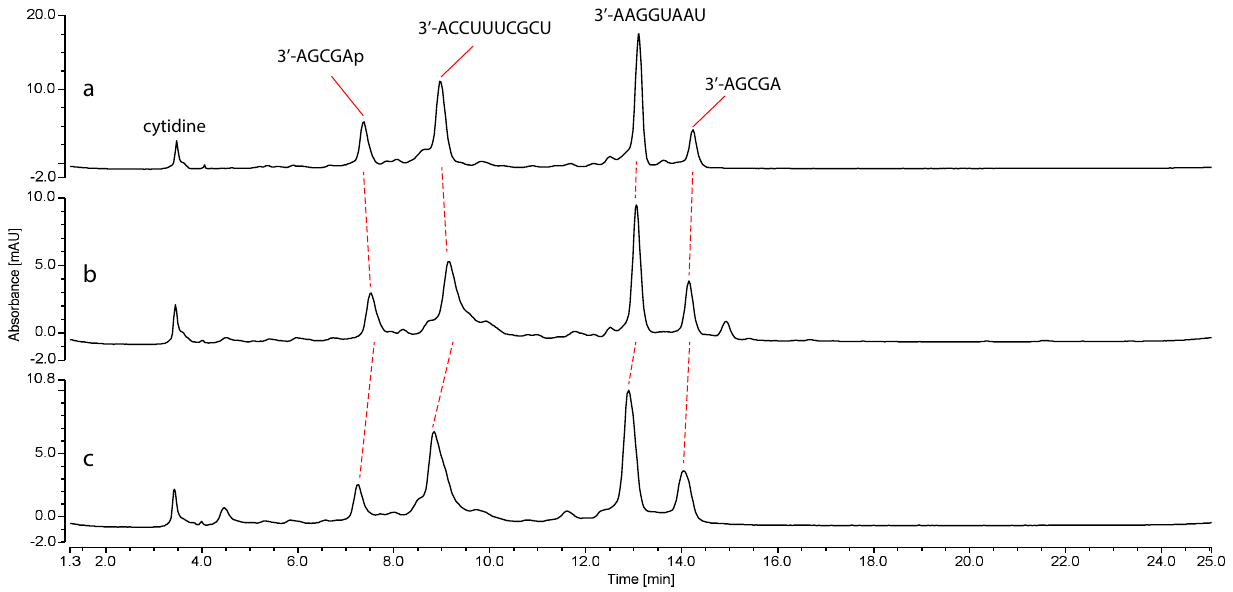

##### Figure SI-18. HPLC traces of the control reaction of 5'P-5mer (6) with 10mer (3) and 8mer (8).

Nicked duplex sequence:

3’AGCGAp

5’UCGCUUUCCA

3’AAGGUAAU

Reactions were monitored using HPLC with 260 nm UV detection. The solution was incubated under the desired conditions and aliquots of 2 μL of the reaction solution were diluted in 18 μL water, the diluted solutions were injected into an HPLC. a. Reaction after 0 hours; b. Reaction after 18 hours at room temperature; c. Reaction after 7 days at -16 °C.

#### Phosphoramidate (2) formation

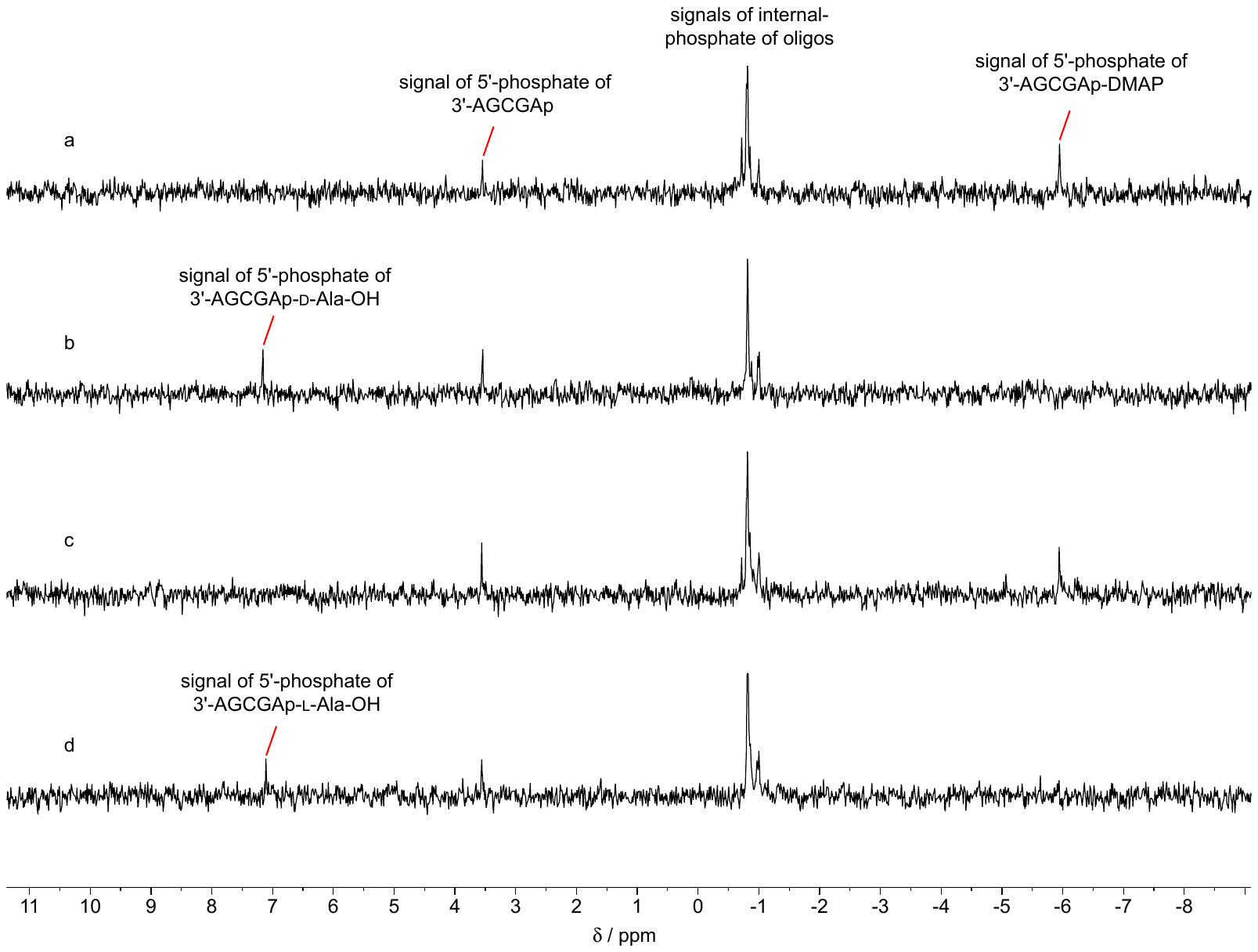

##### Figure SI-19. Stacked ^31^P-NMR spectra for the synthesis of 3'AGCGAp-D-Ala-OH (2-D-Ala) and 3'AGCGAp-L-Ala-OH (2-L-Ala).

The reaction between 3'AGCGAp-DMAP with H-D-Ala-OH in H_2_O/D_2_O solution. a. t = 0 min; b. t = 18 hours. The reaction between 3'AGCGAp-DMAP with H-L-Ala-OH in H_2_O/D_2_O solution. c. t = 0 min; d. t= 18 hours.

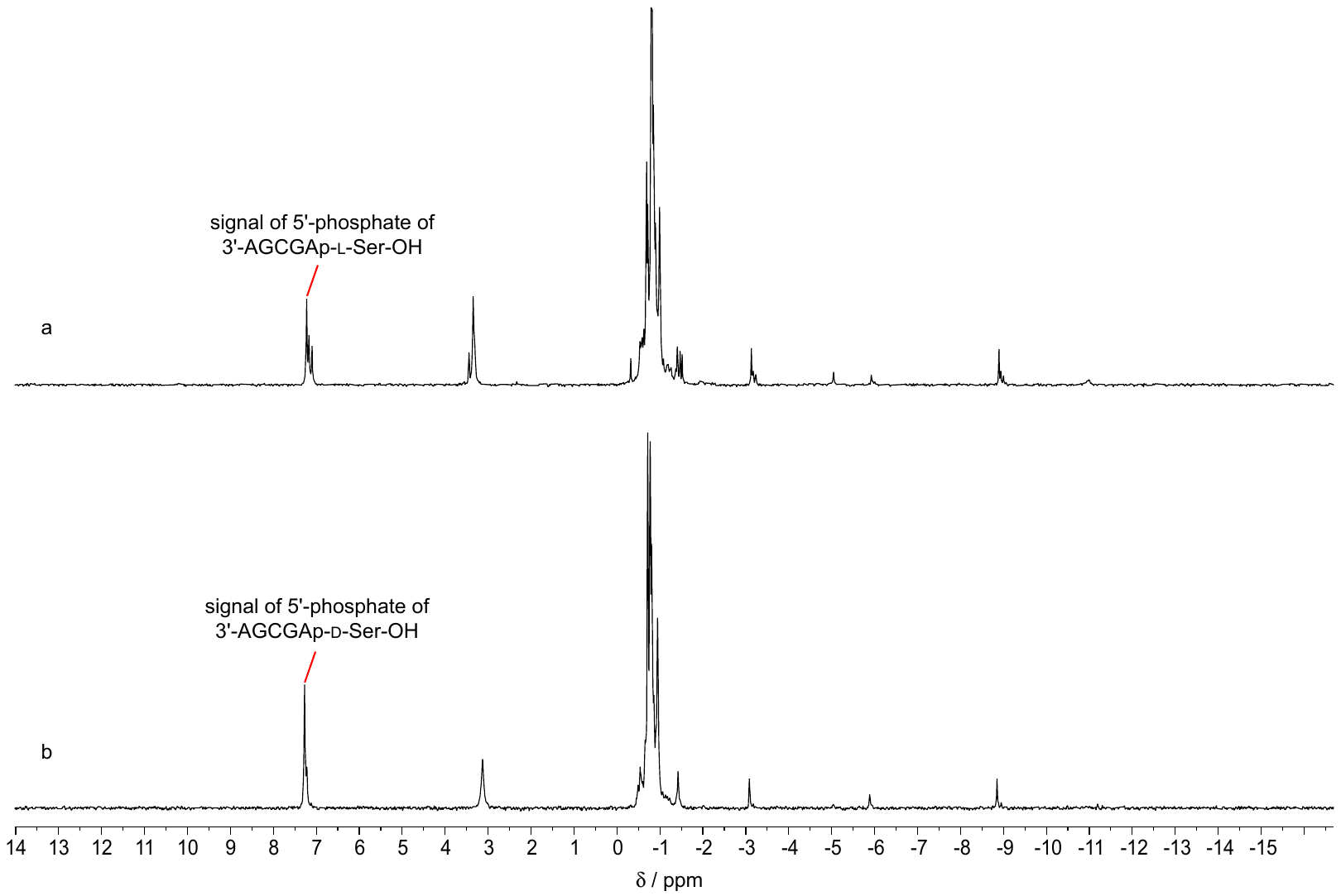

##### Figure SI-20. Stacked ^31^P-NMR spectra for the synthesis of 3'AGCGAp-D-Ser-OH (2-D-Ser) and 3'AGCGAp-L-Ser-OH (2-L-Ser).

a. The reaction between 3'AGCGAp-DMAP with H-L-Ser-OH in H_2_O/D_2_O solution t= 18 hours. b. The reaction between 3'AGCGAp-DMAP with H-D-Ser-OH in H_2_O/D_2_O solution t= 18 hours.

#### Hydrolyses

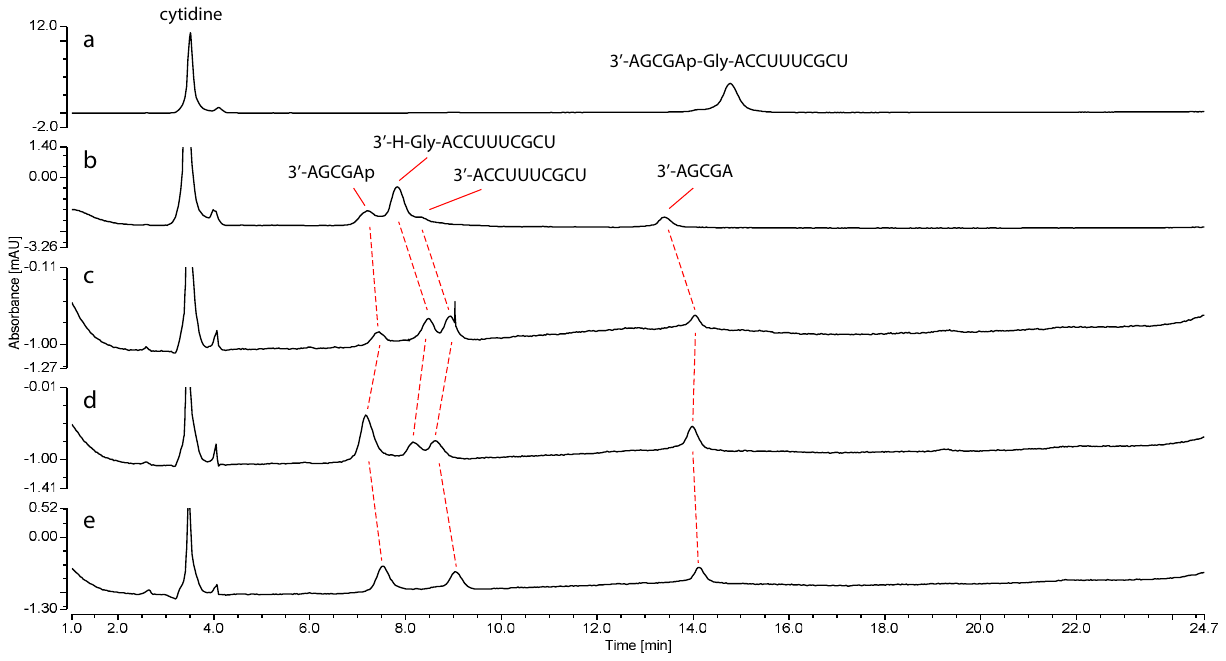

##### Figure SI-21. HPLC traces of the acid hydrolysis of Glycine phosphoramidate-ester (4-Gly)

Reactions were monitored using HPLC with 260 nm UV detection. The solution was incubated at 25 °C in formate buffer (pH 3, 60-83 mM) and aliquots of the reaction solutions were injected into an HPLC. a. hydrolysis of **4-Gly** after 0 hours; b. hydrolysis of **4-Gly** after 17 hours; c. sample b after spiking with 10mer **3**; d. sample c after spiking with 5'P-5mer **6**; e. sample d after base hydrolysis.

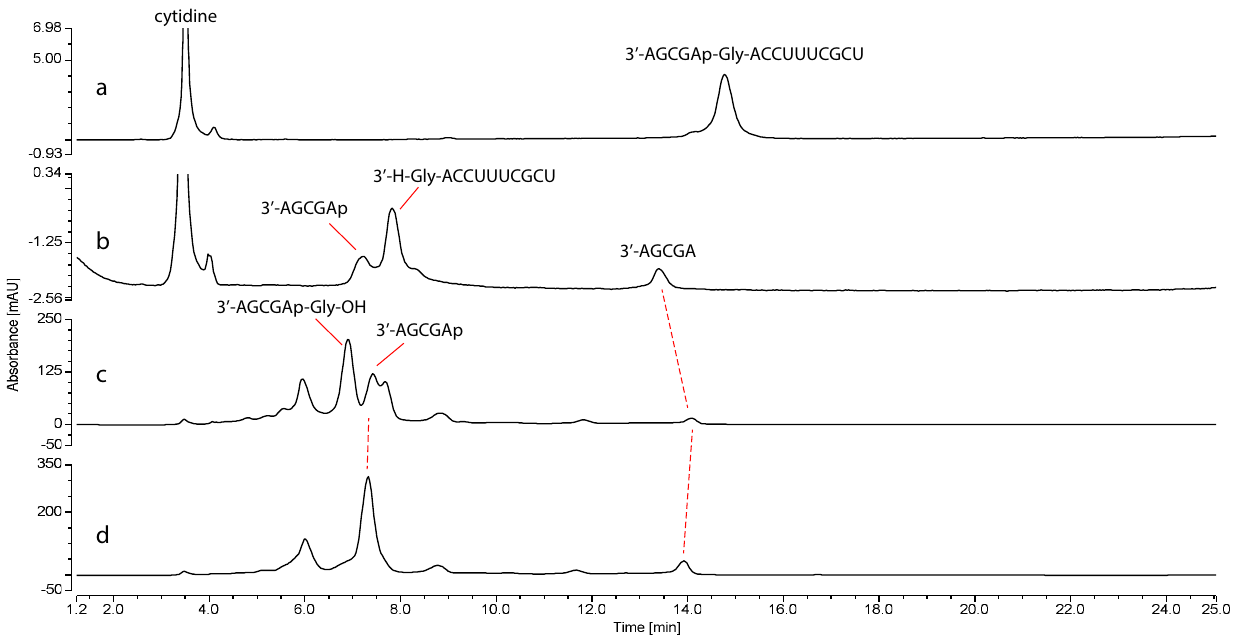

##### Figure SI-22. HPLC traces of the acid hydrolysis of Glycine phosphoramidate-ester (4-Gly) and Glycine amidate-RNA (2-Gly)

Reactions were monitored using HPLC with 260 nm UV detection. The solution was incubated at 25 °C in formate buffer (pH 3, 60-83 mM) and aliquots of the reaction solutions were injected into an HPLC. a. hydrolysis of **4-Gly** after 0 hours; b. hydrolysis of **4-Gly** after 17 hours; c. Hydrolysis of **2-Gly** after 0 hours; b. Hydrolysis of **2-Gly** after 17 hours.

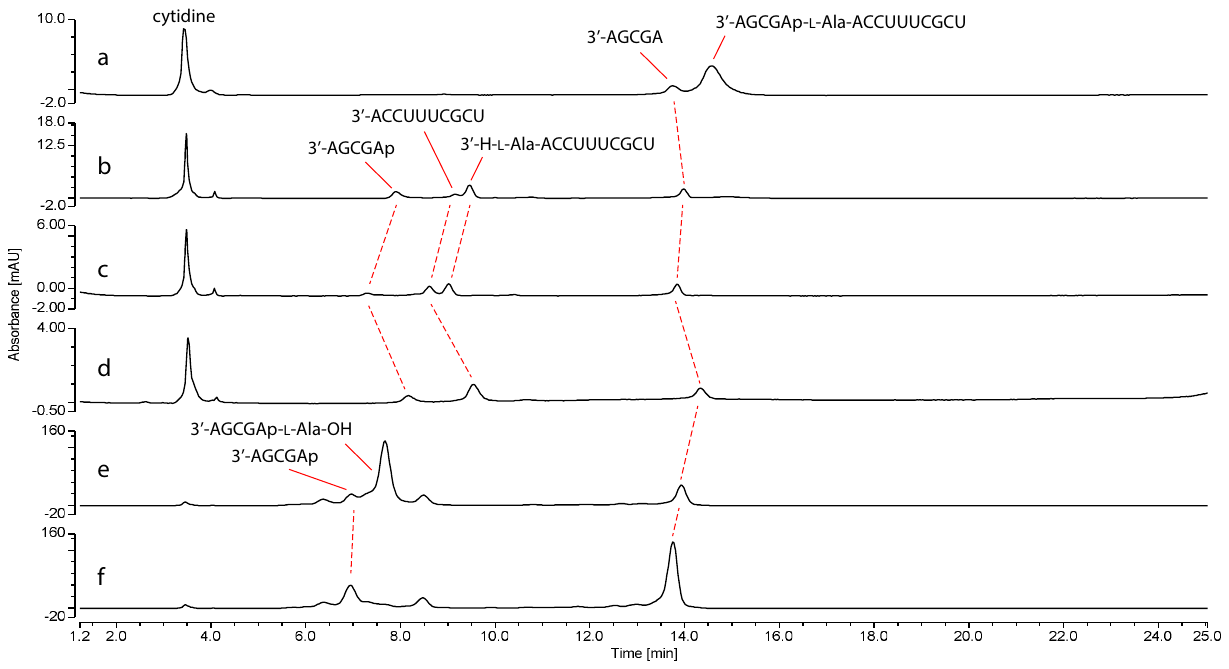

##### Figure SI-23. HPLC traces of the acid hydrolysis of L-Alanine phosphoramidate-ester (4-L-Ala) and L-Alanine amidate-RNA (2-L-Ala)

Reactions were monitored using HPLC with 260 nm UV detection. The solution was incubated at 25 °C in formate buffer (pH 3, 60-83 mM) and aliquots of the reaction solutions were injected into an HPLC. a. hydrolysis of **4-L-Ala** after 0 hours; b. hydrolysis of **4-L-Ala** after 17 hours; c. sample b spiked with 10mer **3**; d. sample b after base hydrolysis; e. Hydrolysis of **2-L-Ala** after 0 hours; f. Hydrolysis of **2-L-Ala** after 17 hours.

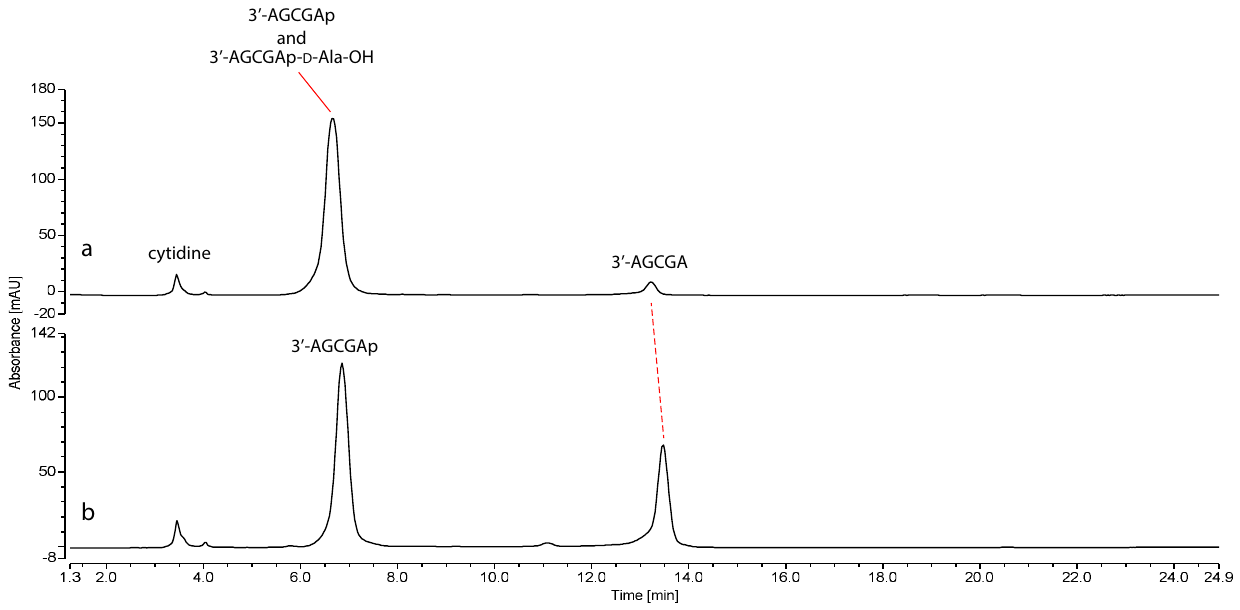

##### Figure SI-24. HPLC traces of the acid hydrolysis of D-Alanine amidate-RNA (2-D-Ala)

Reactions were monitored using HPLC with 260 nm UV detection. The solution was incubated at 25 °C in formate buffer (pH 3, 83 mM) and aliquots of the reaction solutions were injected into an HPLC. a. Hydrolysis of **2-D-Ala** after 0 hours; b. Hydrolysis of **2-D-Ala** after 17 hours.

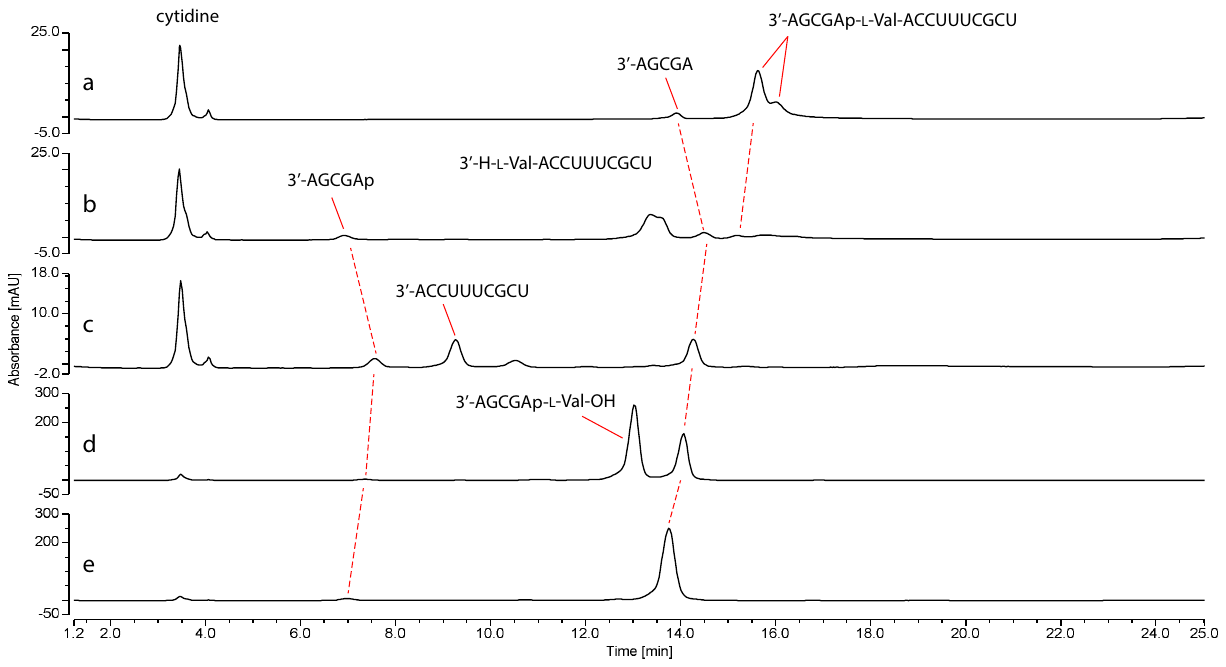

##### Figure SI-25. HPLC traces of the acid hydrolysis of L-valine phosphoramidate-ester (4-L-Val) and L-valine amidate-RNA (2-L-Val)

Reactions were monitored using HPLC with 260 nm UV detection. The solution was incubated at 25 °C in formate buffer (pH 3, 60-83 mM) and aliquots of the reaction solutions were injected into an HPLC. a. hydrolysis of **4-L-Val** after 0 hours; b. hydrolysis of **4-L-Val** after 17 hours; c. sample b after base hydrolysis; e. Hydrolysis of **2-L-Val** after 0 hours; f. Hydrolysis of **2-L-Val** after 17 hours.

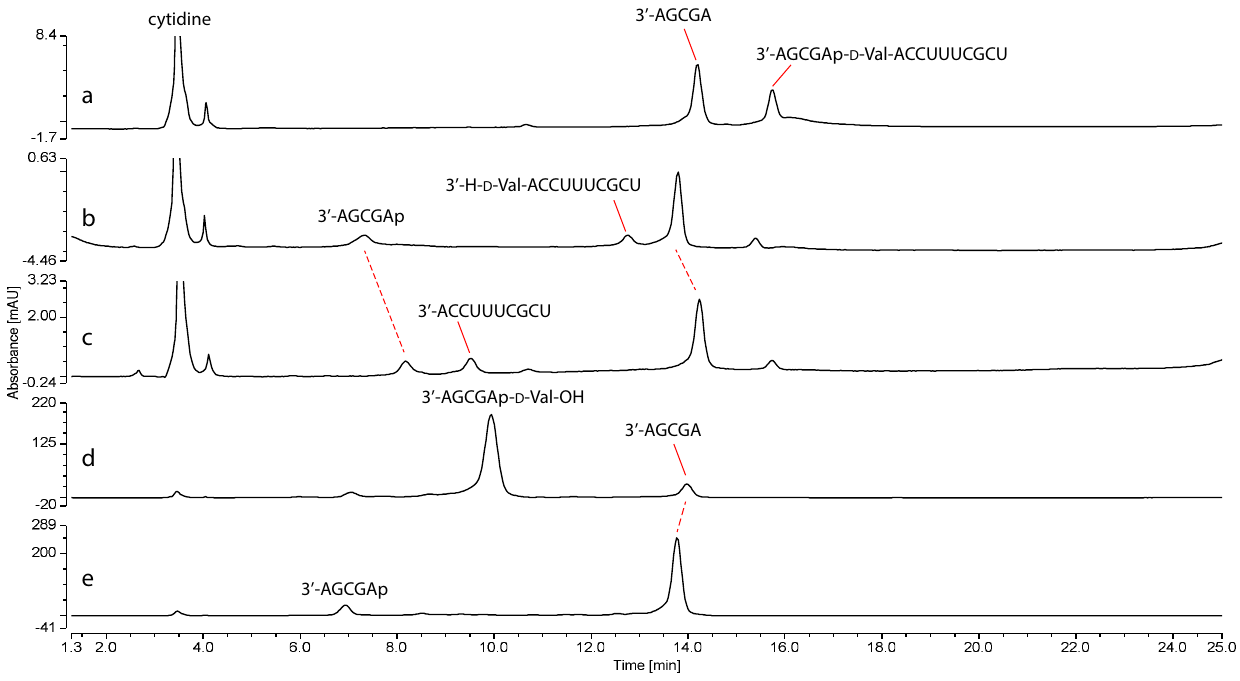

##### Figure SI-26. HPLC traces of the acid hydrolysis of D-valine phosphoramidate-ester (4-D-Val) and D-valine amidate-RNA (2-D-Val)

Reactions were monitored using HPLC with 260 nm UV detection. The solution was incubated at 25 °C in formate buffer (pH 3, 60-83 mM) and aliquots of the reaction solutions were injected into an HPLC. a. hydrolysis of **4-D-Val** after 0 hours; b. hydrolysis of **4-D-Val** after 17 hours; c. sample b after base hydrolysis; e. Hydrolysis of **2-D-Val** after 0 hours; f. Hydrolysis of **2-D-Val** after 17 hours.

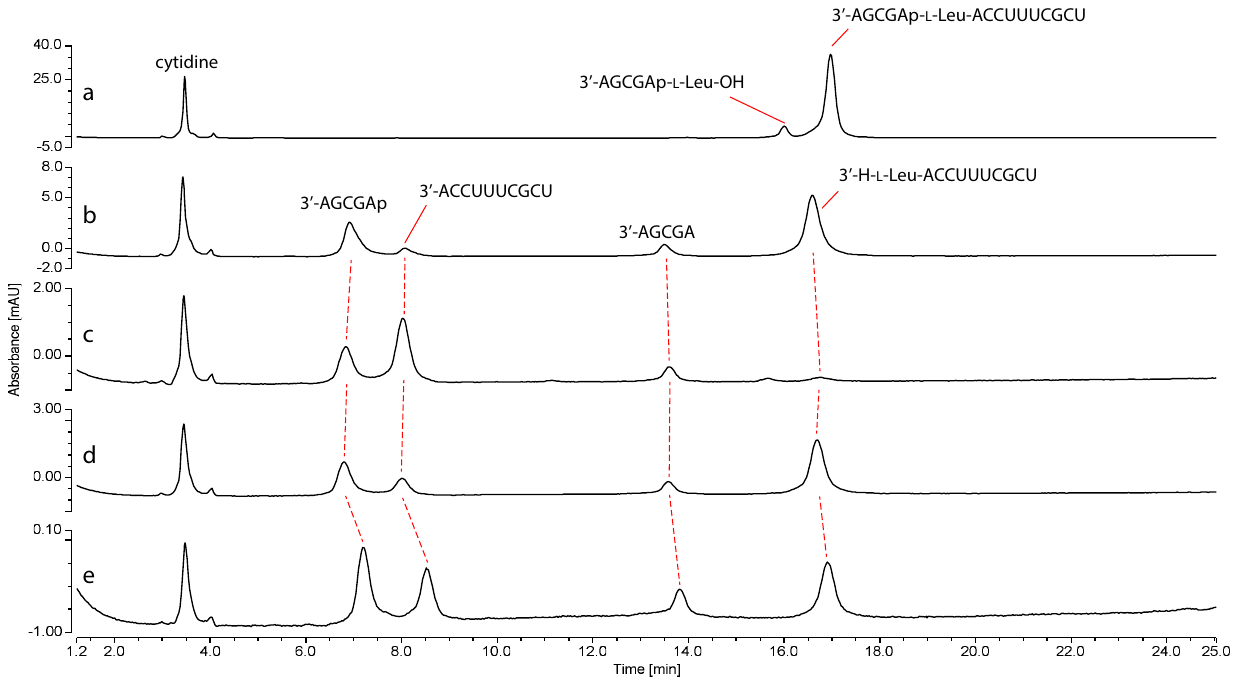

##### Figure SI-27. HPLC traces of the acid hydrolysis of L-Leucine phosphoramidate-ester RNA ester (4-L-Leu)

Reactions were monitored using HPLC with 260 nm UV detection. The solution was incubated at 25 °C in formate buffer (pH 3, 60-83 mM) and aliquots of the reaction solutions were injected into an HPLC. a. hydrolysis of **4-L-Leu** after 0 hours; b. hydrolysis of **4-L-Leu** after 17 hours; c. sample b after base hydrolysis; d. sample b after spiking with 10mer **3**; e. sample d after spiking with 5'P-5mer **6**.

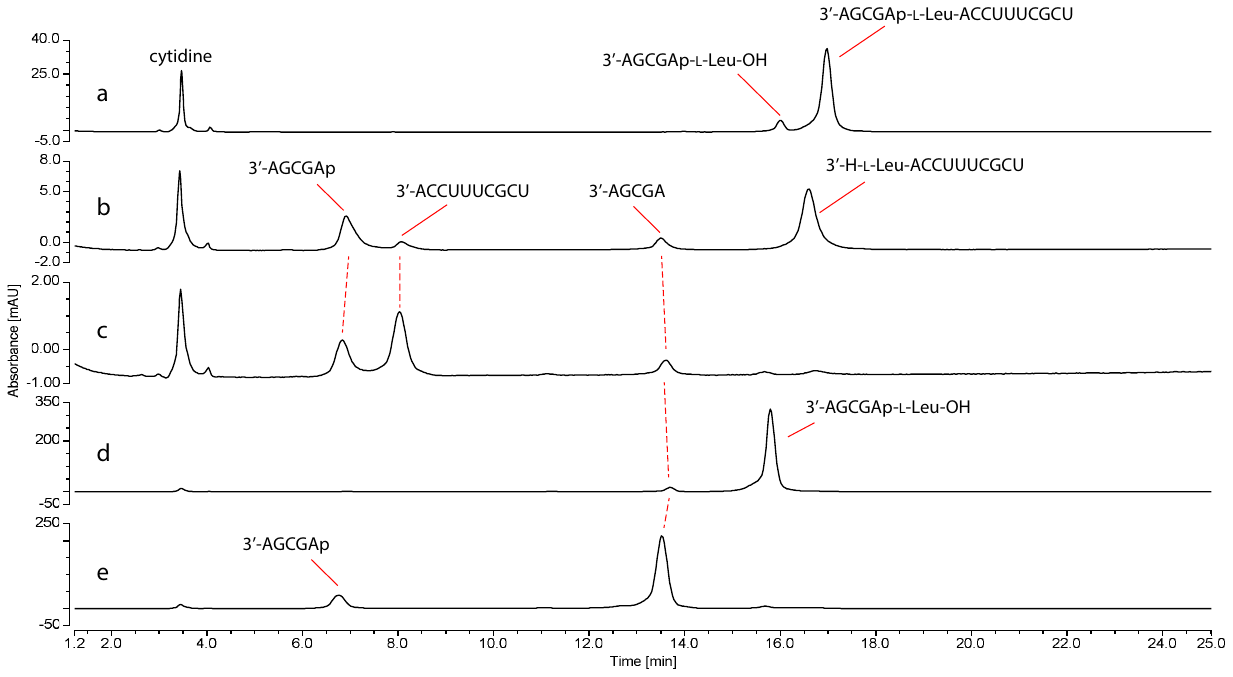

##### Figure SI-28. HPLC traces of the acid hydrolysis of L-Leucine phosphoramidate-ester (4-L-Leu) and L-Leucine amidate-RNA (2-L-Leu)

Reactions were monitored using HPLC with 260 nm UV detection. The solution was incubated at 25 °C in formate buffer (pH 3, 60-83 mM) and aliquots of the reaction solutions were injected into an HPLC. a. hydrolysis of **4-L-Leu** after 0 hours; b. hydrolysis of **4-L-Leu** after 17 hours; c. sample b after base hydrolysis; d. Hydrolysis of **2-L-Leu** after 0 hours; e. Hydrolysis of **2-L-Leu** after 17 hours.

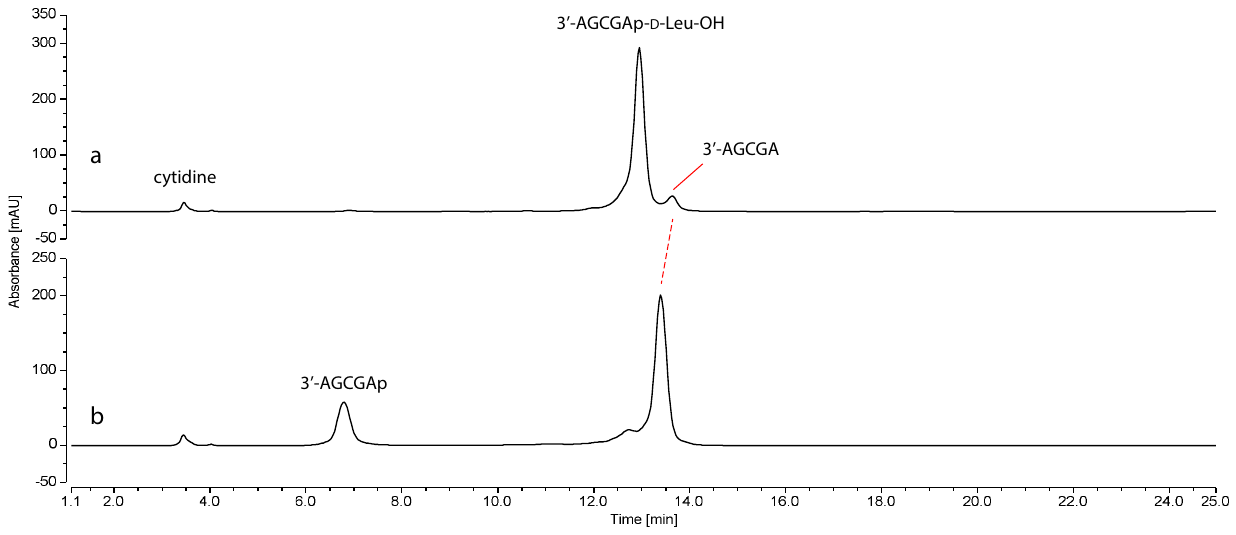

##### Figure SI-29. HPLC traces of the acid hydrolysis of D-Leucine amidate-RNA (2-D-Leu)

Reactions were monitored using HPLC with 260 nm UV detection. The solution was incubated at 25 °C in formate buffer (pH 3, 83 mM) and aliquots of the reaction solutions were injected into an HPLC. a. Hydrolysis of **2-D-Leu** after 0 hours; b. Hydrolysis of **2-D-Leu** after 17 hours.

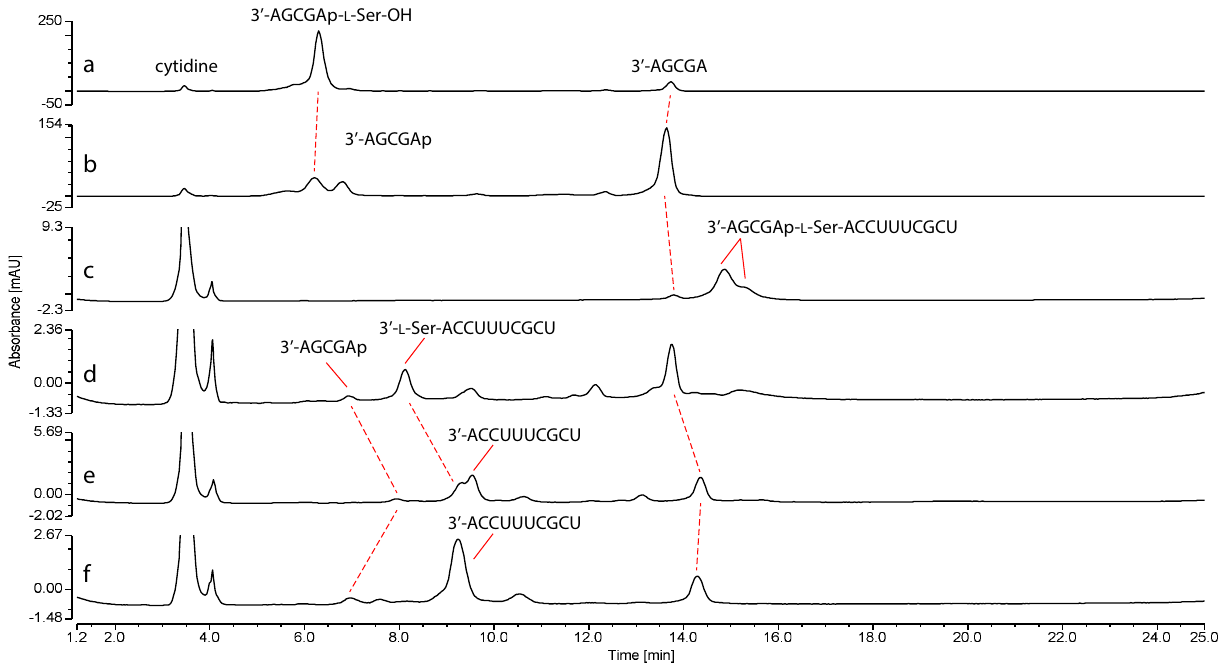

##### Figure SI-30. HPLC traces of the acid hydrolysis of L-Serine phosphoramidate-ester (4-L-Ser) and L-Serine amidate-RNA (2-L-Ser)

Reactions were monitored using HPLC with 260 nm UV detection. The solution was incubated at 25 °C in formate buffer (pH 3, 60-83 mM) and aliquots of the reaction solutions were injected into an HPLC. a. Hydrolysis of **2-L-Ser** after 0 hours; b. Hydrolysis of **2-L-Ser** after 17 hours; c. hydrolysis of **4-L-Ser** after 0 hours; d. hydrolysis of **4-L-Ser** after 17 hours; e. sample d after spiking with 10mer **3**; f. sample e after base hydrolysis.

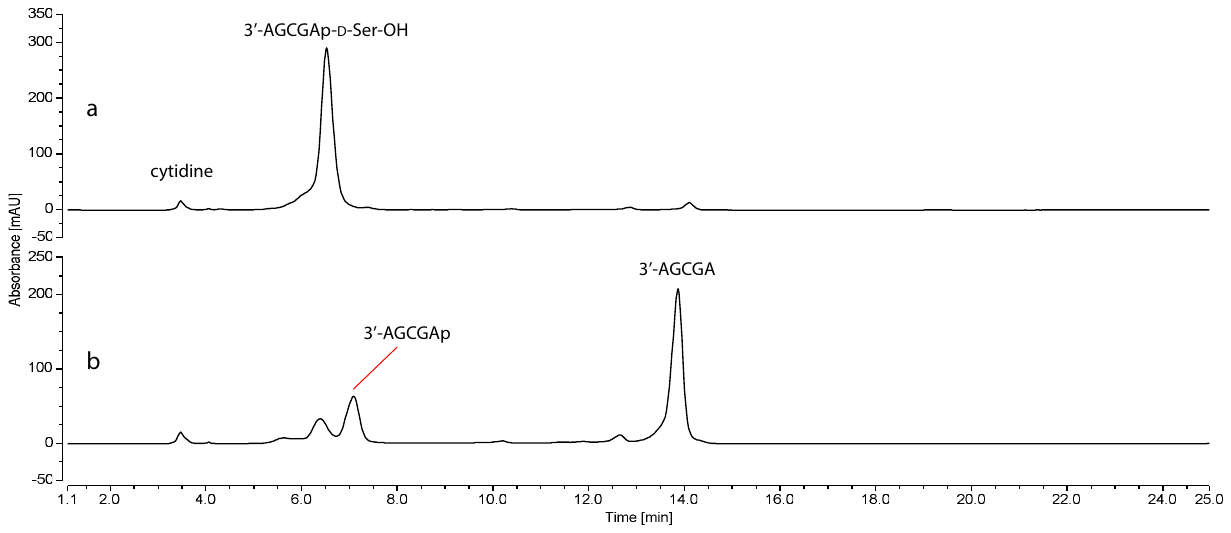

##### Figure SI-31. HPLC traces of the acid hydrolysis of D-Serine amidate-RNA (2-D-Ser)

Reactions were monitored using HPLC with 260 nm UV detection. The solution was incubated at 25 °C in formate buffer (pH 3, 83 mM) and aliquots of the reaction solutions were injected into an HPLC. a. Hydrolysis of **2-D-Ser** after 0 hours; b. Hydrolysis of **2-D-Ser** after 17 hours.

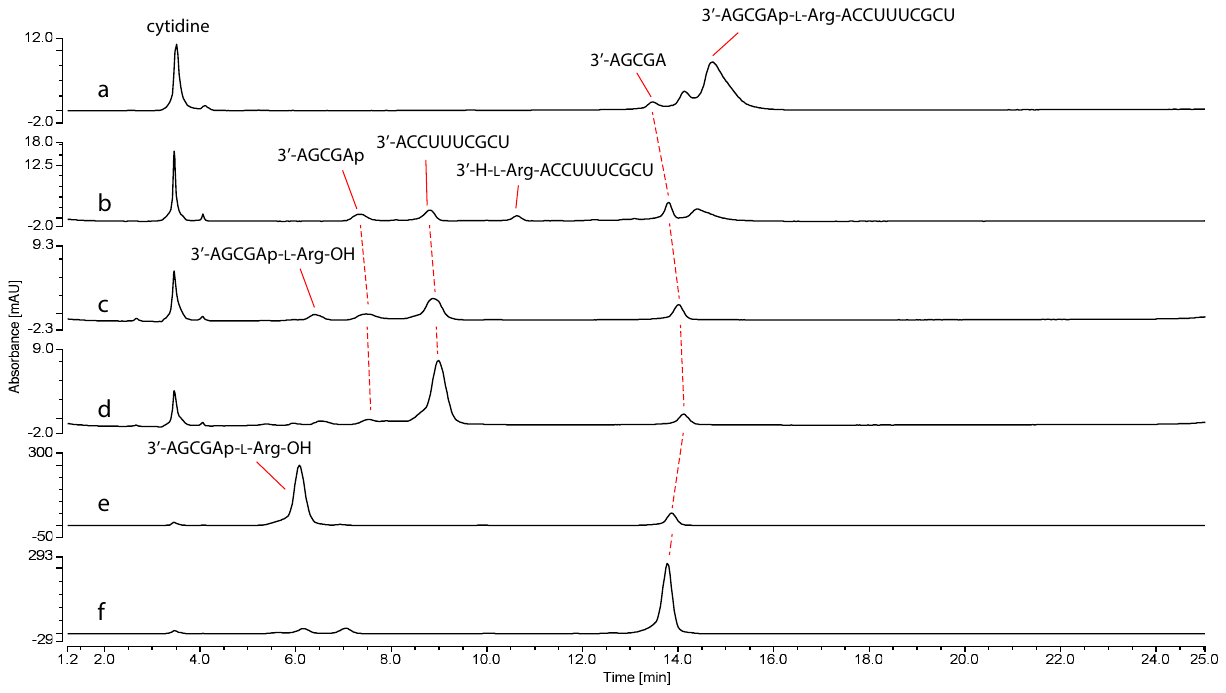

##### Figure SI-32. HPLC traces of the acid hydrolysis of L-Arginine phosphoramidate-ester (4-L-Arg) and L-Arginine amidate-RNA (2-L-Arg)

Reactions were monitored using HPLC with 260 nm UV detection. The solution was incubated at 25 °C in formate buffer (pH 3, 60-83 mM) and aliquots of the reaction solutions were injected into an HPLC. a. hydrolysis of **4-L-Arg** after 0 hours; b. hydrolysis of **4-L-Arg** after 17 hours; c. sample b after base hydrolysis; d. sample c after spiking with 10mer **3**; e. hydrolysis of **2-L-Arg** after 0 hours; f. hydrolysis of **2-L-Arg** after 17 hours.

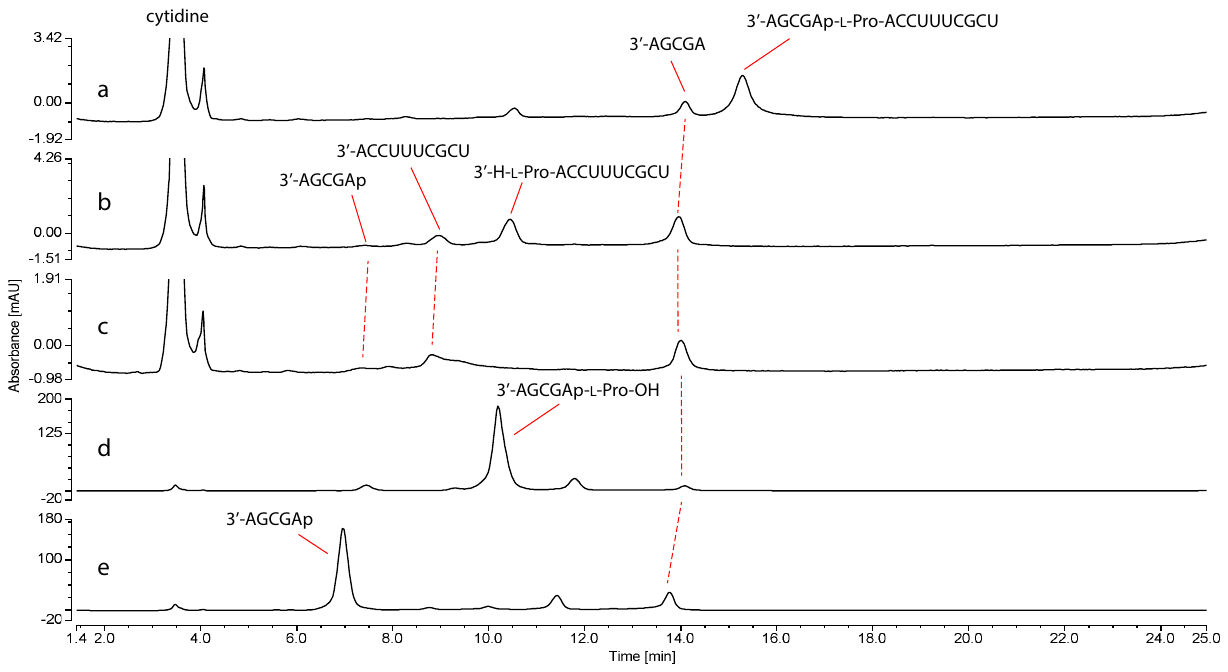

##### Figure SI-33. HPLC traces of the acid hydrolysis of L-Proline phosphoramidate-ester (4-L-Pro) and L-Arginine amidate-RNA (2-L-Pro)

Reactions were monitored using HPLC with 260 nm UV detection. The solution was incubated at 25 °C in formate buffer (pH 3, 60-83 mM) and aliquots of the reaction solutions were injected into an HPLC. a. hydrolysis of **4-L-Pro** after 0 hours; b. hydrolysis of **4-L-Pro** after 17 hours; c. sample b after base hydrolysis; d. hydrolysis of **2-L-Pro** after 0 hours; e. hydrolysis of **2-L-Pro** after 17 hours.

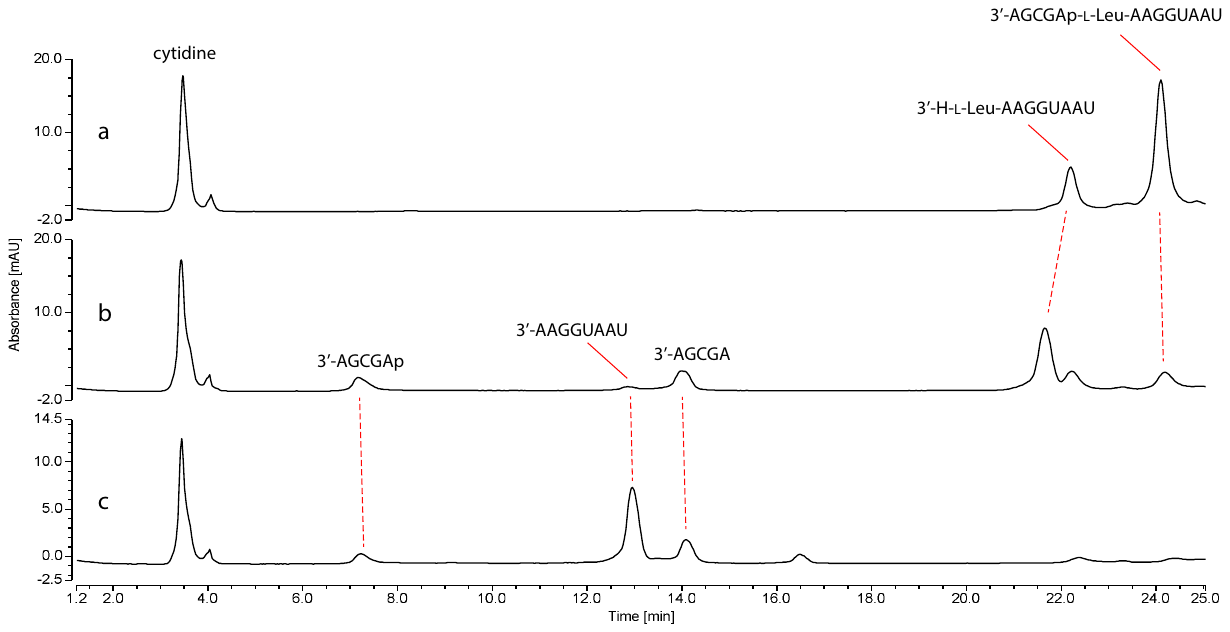

##### Figure SI-34. HPLC traces of the acid hydrolysis of nicked duplex RNA-L-Leucine phosphoramidate-ester (7-L-Leu).

Reactions were monitored using HPLC with 260 nm UV detection. The solution was incubated at 25 °C in formate buffer (pH 3, 60-83 mM) and aliquots of the reaction solutions were injected into an HPLC. a. hydrolysis of **7-L-Leu** after 0 hours; b. hydrolysis of **7-L-Leu** after 17 hours; c. sample b after base hydrolysis.

### Additional Data

#### Control for Reaction of Free Amino Acid Under Phosphoramidate Ester Forming Conditions

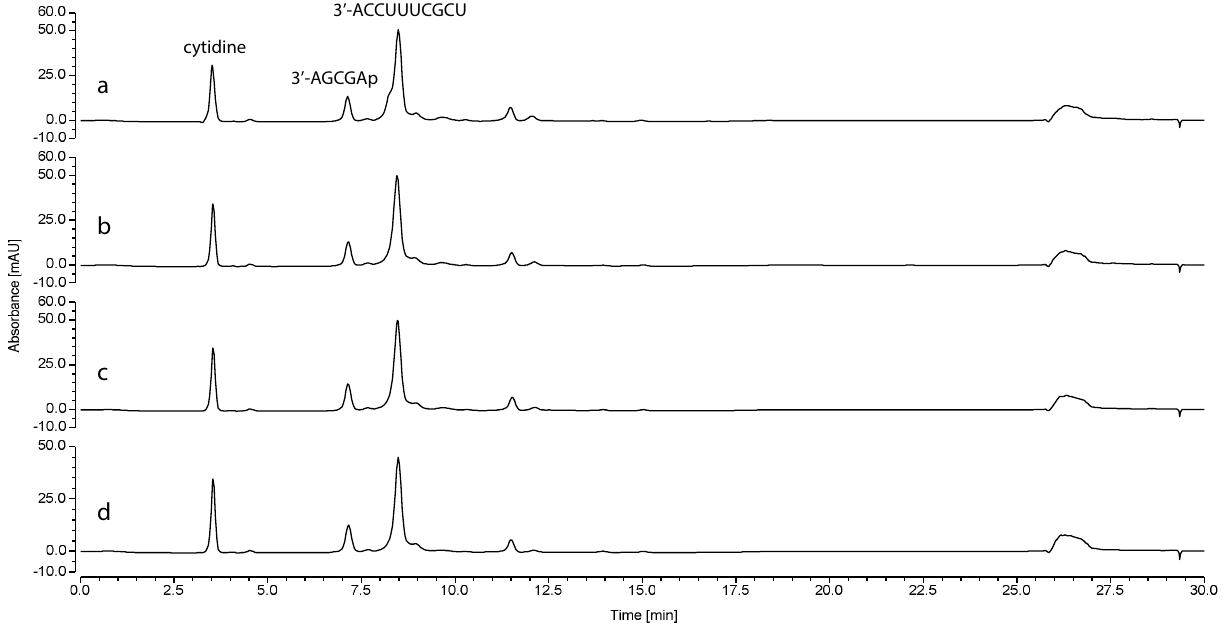

##### Figure SI-35. HPLC traces of the lack of formation of RNA-aminoacyl- phosphoramidate-ester (4) in the absence of (2).

Loop duplex sequence:

3’AGCGAp

5’UCGCUUUCCA

Reactions were monitored using HPLC with 260 nm UV detection. The solution was incubated at 20 °C for 18 hours. After the desired time each aliquot was diluted in 18 μL water, the diluted solutions were injected into an HPLC. a. without any amino acid addition; b. with Gly (10 mM); c. with L-Ala (10 mM); d. with D-Ala (10 mM).

#### Phosphoramidate (2) and Phosphoramidate-Ester (4) Mass Spec data

##### TABLE SI-1. Summary of MALDI and LCMS identification of phosphoramidates (2) and phosphoramidate-esters (4) or (7). LC traces for 2-Gly, 2-L-Ala, 2-D-Val and 2-D-Ser provided below (Figure SI-35 to SI-38).

| Amino Acid | Phosphoramidate (2) | | Phosphoramidate-ester (4 or 7) | |
| --- | --- | --- | --- | --- |
|  | Calculated [M+H] | Found | Calculated  [M+H] | Found |
| Glycine | 1729.3 | LCMS: 863.0 [(M-2H)/2] | 4767.7 | MALDI: 4768.0 [M+H] |
| L-Ala | 1743.3 | LCMS: 869.9 [(M-2H)/2] | 4781.7 | MALDI: 4782.2 [M+H] |
| D-Ala | 1743.3 | MALDI: 1765.1 [M+Na] | - | - |
| L-Val | 1771.3 | MALDI: 1770.7 [M+H] | 4809.7 | MALDI: 4810.0 [M+H] |
| D-Val | 1771.3 | LCMS: 884.0 [(M-H)/2] | - | - |
| L-Leu | 1785.3 | MALDI: 1785.6 [M+H] | 4823.7 | MALDI: 4823.8 [M+H] |
| D-Leu | 1785.3 | MALDI: 891.1 [M+H] | - | - |
| L-Ser | 1759.3 | MALDI: 1759.1 [M+H] | 4797.7 | MALDI: 4798.2 [M+H] |
| D-Ser | 1759.3 | LCMS: 877.9 [(M-H)/2] | - | - |
| L-Arg | 1828.4 | MALDI: 1828.5 [M+H] | 4866.8 | MALDI: 4866.6 [M+H] |
| L-Pro | 1769.3 | MALDI: 1769.2 [M+H] | 4807.7 | MALDI: 4982.8 [M+8Na] |
| Nicked Duplex | - | - | 4323.7 | MALDI: 4324.2 [M+H] |

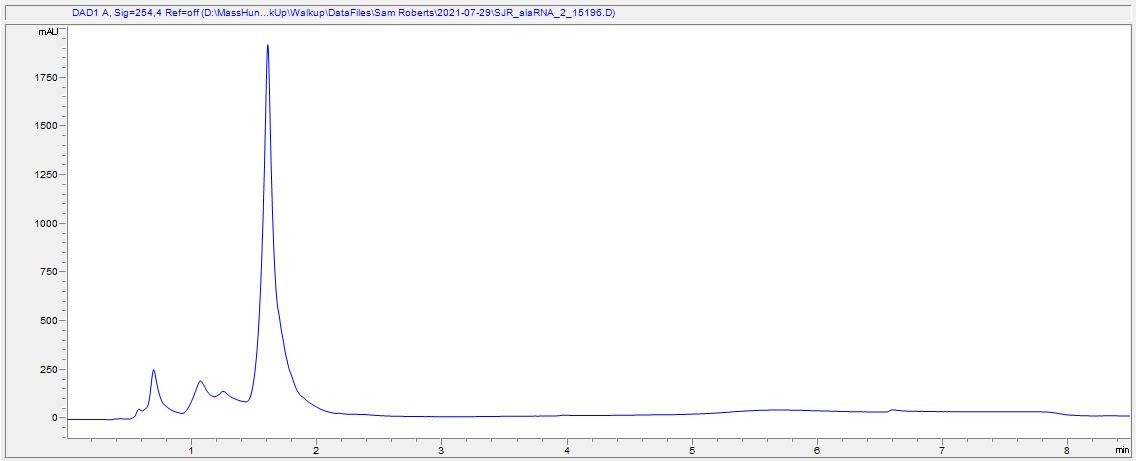

##### Figure SI-36. LC traces L-Alanine phosphoramidate-ester (2-L-Ala).

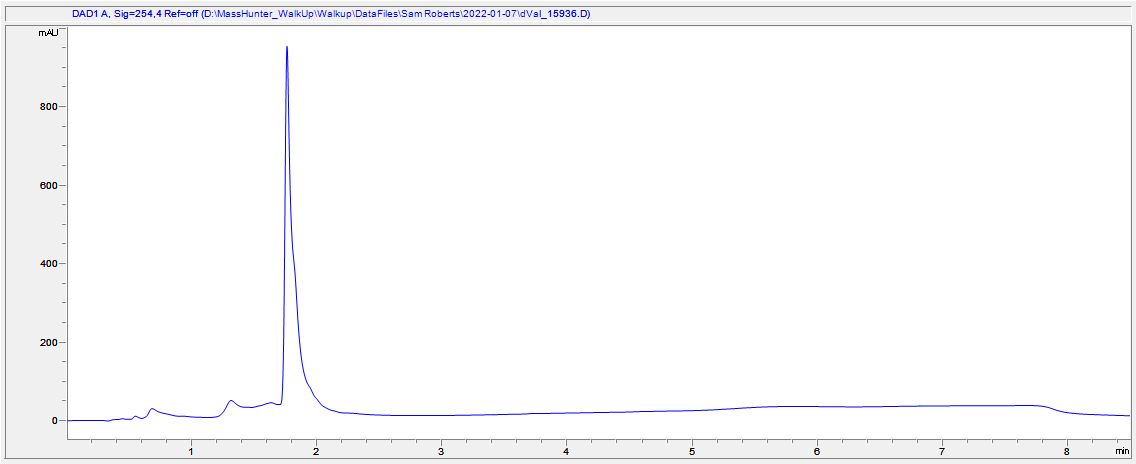

##### Figure SI-37. LC traces D-Valine phosphoramidate-ester (2-D-Val).

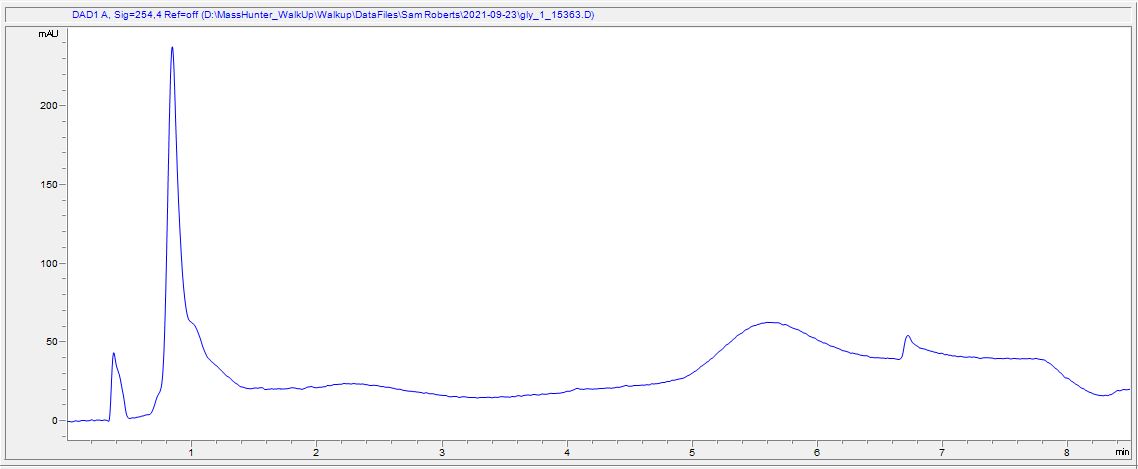

##### Figure SI-38. LC traces Glycine phosphoramidate-ester (2-Gly).

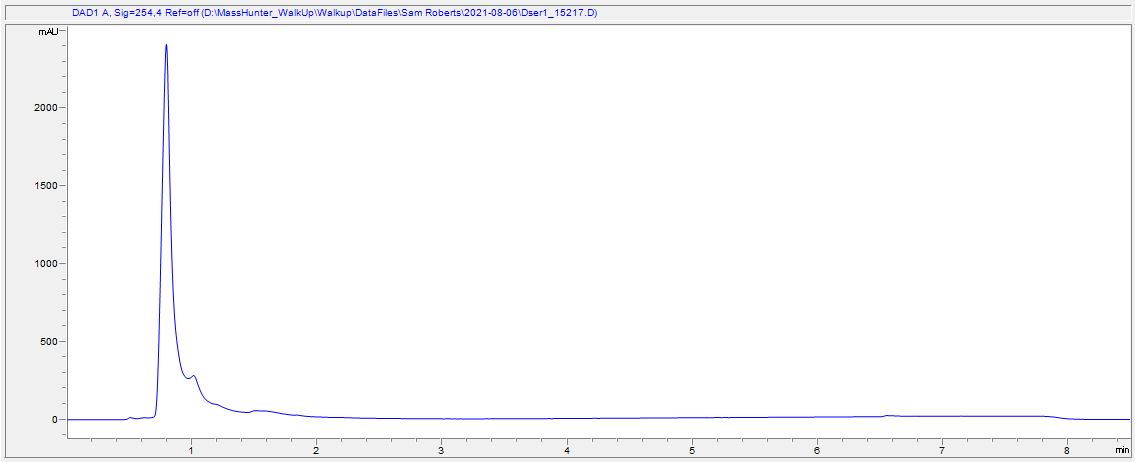

##### Figure SI-39. LC traces D-Serine phosphoramidate-ester (2-D-Ser).

#### Yield of 5'P-5mer (6) produced from hydrolysis of phosphoramidate RNA (2)

##### TABLE SI-2. Yield of 5'P-5mer (6) produced from consumption of phosphoramidate RNA (2) by hydrolysis in pH 3 formate buffer

| Amino Acid | % Yield (6) | % Yield (9) | Amino Acid | % Yield (6) | % Yield (9) |
| --- | --- | --- | --- | --- | --- |
| Gly | 85% | 15% | - | - | - |
| L-Ala | 24% | 76% | D-Ala | 24% | 76% |
| L-Leu | 16% | 84% | D-Leu | 16% | 84% |
| L-Val | 3% | 97% | D-Val | 3% | 97% |
| L-Ser | 15% | 85% | D-Ser | 15% | 85% |
| L-Arg | 6% | 94% | L-Pro | 6% | 93% |

#### HPLC Calibration Curves

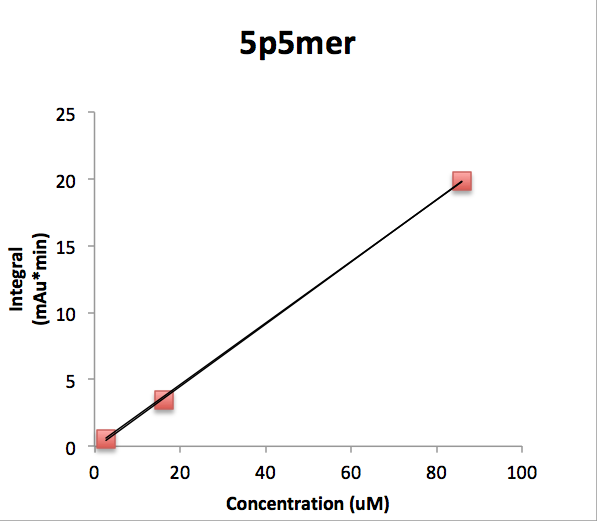

##### CHART SI-1. Calibration curve of 5'P-5mer (6) for quantification of yields by HPLC. y=kx, k = 0.2304 mAu•min•μM^-1^. R^2^= 0.99967.

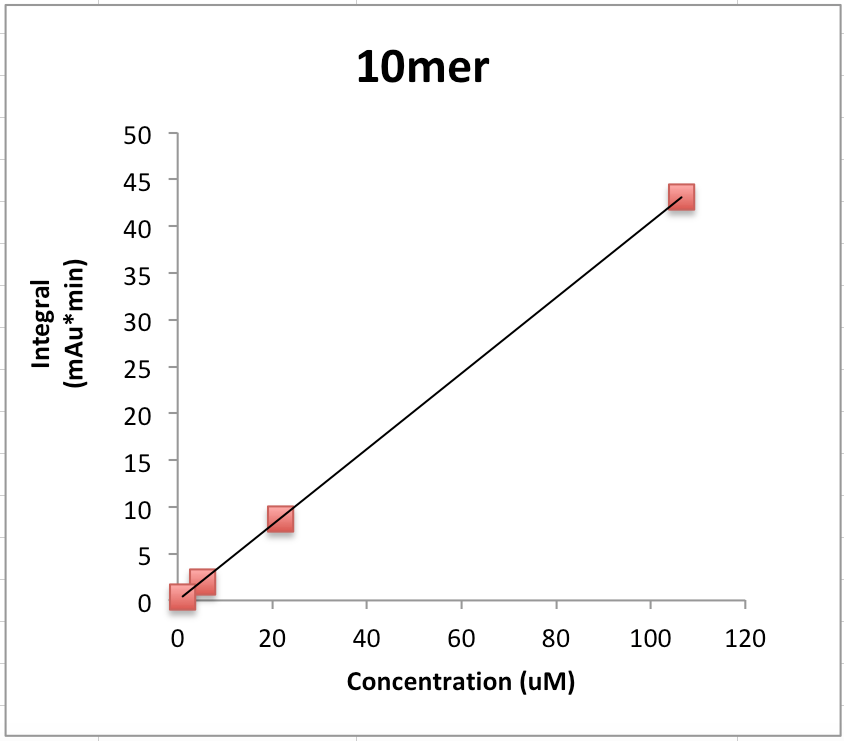

##### CHART SI-2. Calibration curve of 10mer (3) for quantification of yields by HPLC. y=kx, k = 0.4046 mAu•min•μM^-1^. R^2^= 0.99996.

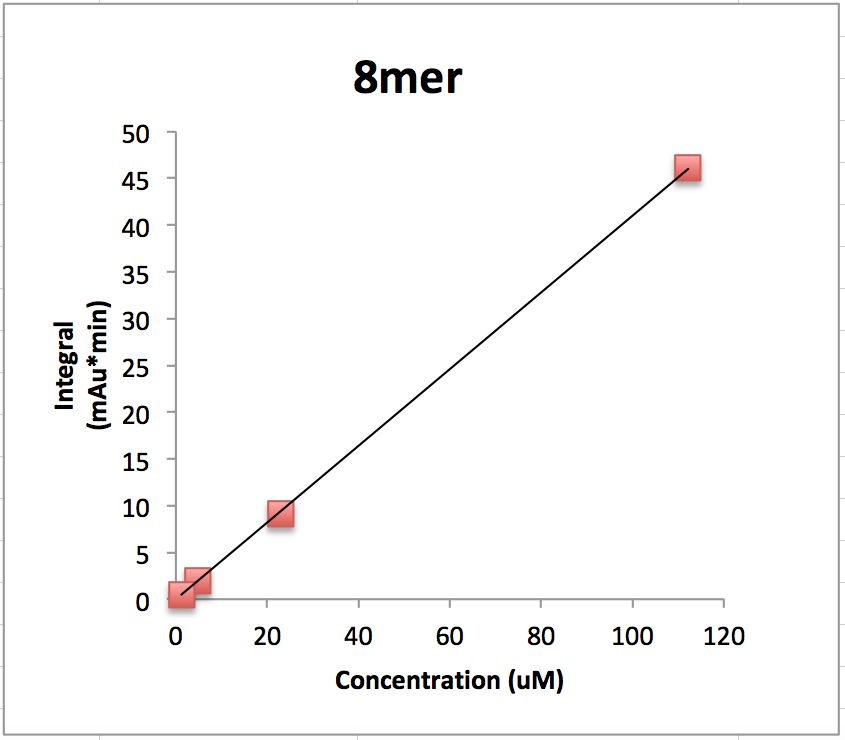

##### CHART SI-3. Calibration curve of 8mer (8) for quantification of yields by HPLC. y=kx, k = 0.4100 mAu•min•μM^-1^. R^2^= 0.99992.

#### Stereoselectivity of phosphoramidate-ester (4) formation

##### TABLE SI-3. Ratio of yields of formation L:D phosphoramidate-esters (4). Quantified as the average of three replicate ratios.

| Amino Acid | Room temperature 18 hours | -16 °C for 7 days | -16 °C for 14 days |
| --- | --- | --- | --- |
| Ala | 9.8:1 | 18.4:1 | 9.9:1 |
| Leu | 11.6:1 | 37.2:1 | 27.6:1 |
| Val | 6.5:1 | 48.1:1 | 10.9:1 |
| Ser | 4.6:1 | 12.0:1 | 10.5:1 |
| Nicked Duplex-Leu | 6.9:1 | 11.7:1 | 29.0:1 |
